# Psilocybin prevents activity-based anorexia in female rats by enhancing cognitive flexibility: contributions from 5-HT1A and 5-HT2A receptor mechanisms

**DOI:** 10.1101/2023.12.12.571374

**Authors:** K Conn, LK Milton, K Huang, H Munguba, J Ruuska, MB Lemus, E Greaves, J Homman-Ludiye, BJ Oldfield, CJ Foldi

## Abstract

Psilocybin has shown promise for alleviating symptoms of depression and is currently in clinical trials for the treatment of anorexia nervosa (AN), a condition that is characterised by persistent cognitive inflexibility. Considering that enhanced cognitive flexibility after psilocybin treatment is reported to occur in individuals with depression, it is plausible that psilocybin could improve symptoms of AN by breaking down cognitive inflexibility. A mechanistic understanding of the actions of psilocybin is required to tailor the clinical application of psilocybin to individuals most likely to respond with positive outcomes. This can only be achieved using incisive neurobiological approaches in animal models. Here, we use the activity-based anorexia (ABA) rat model and comprehensively assess aspects of reinforcement learning to show that psilocybin (post-acutely) improves body weight maintenance in female rats and facilitates cognitive flexibility, specifically via improved adaptation to the initial reversal of reward contingencies. Further, we reveal the involvement of signalling through the serotonin (5-HT) 1A and 5-HT2A receptor subtypes in specific aspects of learning, demonstrating that 5-HT1A antagonism negates the cognitive enhancing effects of psilocybin. Moreover, we show that psilocybin elicits a transient increase and decrease in cortical transcription of these receptors (*Htr2a* and *Htr1a*, respectively), and a further reduction in the abundance of *Htr2a* transcripts in rats exposed to the ABA model. Together, these findings support the hypothesis that psilocybin could ameliorate cognitive inflexibility in the context of AN and highlight a need to better understand the therapeutic mechanisms independent of 5-HT2A receptor binding.

## Introduction

AN is characterised by pathological weight loss driven by restrictive feeding and excessive exercise behaviours, has the highest mortality rate of any psychiatric disorder ^1^, and is the leading cause of death in females aged 15-24 ^2^. Life time prevalence rates of AN are estimated at up to 4% in females and 0.5% in males ^3^ and while selective serotonin reuptake inhibitors (SSRIs) are the leading pharmacological treatment, they do not improve clinical symptoms in underweight individuals with AN ^4^.Cognitive inflexibility may be a trait marker of vulnerability to AN, considering that dysfunction arises before the onset of symptoms ^5^ and persists after weight recovery ^6^. Impairments in cognitive flexibility have been consistently seen in AN patients ^7–10^, and are associated with low quality of life ^11^, making this symptom a primary target for therapeutic intervention. Cognitive flexibility is a fundamental element of executive functioning that allows for behavioural adaptation to a variable environment, and as a consequence, is associated with favourable outcomes throughout the lifespan ^12^. This capability is compromised across a range of neuropsychiatric disorders that include but are not limited to; depression and anxiety disorders, substance-use disorders, obsessive-compulsive disorder and anorexia nervosa (AN) ^13^. In each of these conditions, psilocybin-assisted therapy ^14–17^ has either been shown to elicit positive outcomes or is being currently trialled.

Converging evidence from clinical trials and preclinical studies indicates that psilocybin is an effective treatment for symptoms of several psychiatric disorders ^18^ and may circumvent issues with medication compliance because long-term improvements (at least for depression) have been demonstrated after a single dose ^14^. However, there is little evidence to date that disentangles its pharmacological efficacy from the clinically-guided psychological intervention that accompanies psilocybin exposure in these trials ^19^. Moreover, while the pharmacological actions of psilocybin are now better understood ^20, 21^, how these actions translate to therapeutic outcomes remains unclear. Based on the proposal that the therapeutic effects of psilocybin relate to the promotion of flexible thinking and relaxation of maladaptive, rigidly held beliefs ^14^, and the evidence that psilocybin elicits long lasting effects on cognitive and neural flexibility ^22^, it seems likely that at least some aspects of therapeutic efficacy may be driven by enhanced cognitive flexibility. However, the neurobiological mechanisms through which psilocybin acts to improve cognitive flexibility are unknown, and there are multiple components of learning and cognition that could contribute to enhanced flexible thinking and behaviour after psilocybin treatment that have not been systematically addressed.

There is evidence implicating serotonin (5-HT) dysfunction in AN, with positron emission tomography (PET) imaging studies revealing decreased binding to the 5-HT2A receptor (5-HT2AR) subtype ^23^ and increased binding to the 5-HT1A receptor (5-HT1AR) subtype ^24^ in the frontal cortex of patients. Psilocybin is an agonist for both receptor subtypes ^25^, raising the intriguing possibility that psilocybin could rescue or reverse cognitive inflexibility by re-establishing the balance of 5-HT signalling in those with AN. Whether or not psilocybin has therapeutic effects in individuals with AN will be revealed by ongoing clinical trials (e.g., NCT04052568, NCT04661514, NCT05481736, NCT04505189). However, these trials are not capable of testing the mechanisms through which psilocybin acts to elicit improvements in symptoms; moreover, they have been criticised in recent years for methodological constraints most notably their inability to blind participants to treatment conditions, which can bias outcomes in line with expectancy effects ^26, 27^.

Preclinical studies in animal models are critical for advancing the understanding of the behavioural and pharmacological mechanisms underlying the therapeutic effects of psilocybin ^28^, with evidence converging on increased neuroplasticity as a key driver of beneficial outcomes ^29, 30^. Unfortunately, efforts in this space have focused largely on traditional assays of depression-related behaviour in rodents ^31^, with variable findings of either improvements ^32^ or no effects ^33^, dependent on the assay or animal model used ^34^. Given the growing appreciation in behavioural neuroscience that these types of behavioural tests (i.e. the forced swim test) do not reliably translate to human depressive syndromes ^35^ and that reinforcement learning tasks offer key advantages including more relevant clinical links and repeatability ^36^, these early approaches clearly need to be redressed. Other key methodological details in prior studies need to be considered, particularly the role of multiple dosing (cross-over) designs, antagonising 5-HT2AR with ketanserin (a compound with many known non-serotonergic binding sites ^37^), and the measurable motoric side effects of acute psilocybin administration ^38^.

The investigation of neurochemical or neural circuit substrates of these effects centre around the actions of psilocybin on the serotonin-2 (5-HT2) receptor subtypes ^32, 39–43^ but the evidence for the role of 5-HT2AR in rodent cognitive flexibility is conflicting, where acute activation either impairs ^44^ or has no effect on performance ^45^. Less attention has been paid to the possibility that actions at other 5-HTRs might mediate cognitive effects of psilocybin, despite 5-HT2A *independent* effects seen for alleviation of depression-like behaviour ^32^, dendritic spine formation ^46^, and neuronal synchronicity ^47^. It is likely that specific aspects of psilocybin-induced cognitive flexibility involve other 5-HT receptors ^48, 49^ and their integration with other neuromodulatory systems, most notably dopamine ^50–52^. The challenge in identifying the neuronal substrates for improved flexibility after psilocybin is heightened when attempting to understand whether there may be disorder-specific effects in individuals with AN ^53, 54^, who present with disturbed 5-HT function that remains inadequately understood.

In the present study, our objective was to comprehensively investigate how psilocybin, in a 5-HT receptor-dependent manner, may alter some of the core components that underlie cognitive flexibility, such as incentive motivation and task engagement ^55^, response inhibition ^55^, and reward efficacy ^56^. All animals received psilocybin only once, with or without prior administration of selective 5-HT1A and 5-HT2A receptor antagonists, and learning outcomes were assessed post-acutely. In addition, we used the most well-established rodent model of AN, activity-based anorexia (ABA) ^57^, that elicits rapid body weight loss combined with paradoxical hyperactivity ^58^ to determine whether psilocybin has differential effects on 5-HTR function in the context of eating disorder pathology. ABA rats exhibit impairments in cognitive flexibility on both reversal learning ^59^ and attentional-set shifting tasks ^60^, which is rescued by suppressing cortico-striatal circuitry ^61^ a key site of psychedelic drug action ^62^. Finally, we assessed psilocybin-induced alterations in the abundance of 5-HTR mRNA transcripts in the prefrontal cortex to determine the time-course of effects as well as its impact following the development of the ABA phenotype. Together, these studies reveal specific roles of 5-HT receptor subtypes in enhanced flexible learning after psilocybin and point towards a molecular mechanism that may underpin the efficacy of psilocybin for treating symptoms of AN.

## Methods

### Animals and housing

All animals were obtained from the Monash Animal Research Platform (MARP; Clayton, VIC, Australia). To assess direct effects of psilocybin on the development of the ABA phenotype, female Sprague-Dawley rats (*n*=35 behaviour; *n*=12 RNAscope) were 6 weeks of age on arrival in the laboratory. Young female rats were used in these studies because they are particularly vulnerable to developing the ABA phenotype, a feature that is incompletely understood but has translational relevance to the increased prevalence of AN in young women. In order to asses cognitive and behavioural phenotypes relevant to AN/ABA, we used separate cohorts of aged matched female Sprague-Dawley rats (total *n*=168) that commenced training at 7 weeks of age (see **Supplementary Table 1** for details). To examine the effects of psilocybin on 5-HTR subtype abundance across a time course, an additional cohort (*n*=19) of female Sprague-Dawley rats were used, with administration matched to behavioural cohorts at 8 weeks of age. In all cases, animals were group-housed and acclimated to the 12h light/dark cycle (lights off at 1100h) for 7 days in a temperature (22-24°C) and humidity (30-50%) controlled room before experiments commenced. Because the behavioural aspects of ABA (i.e. wheel running and food intake) as well as aspects of reinforcement learning are known to fluctuate with the oestrous cycle in female rats ^63, 64^, a male rat was individually housed in all experimental rooms at least 7 days prior to experimentation in order to facilitate synchronisation of cycling, known as the Whitten Effect ^65^. All experimental procedures were conducted in accordance with the Australian Code for the care and use of animals for scientific purposes and approved by the Monash Animal Resource Platform Ethics Committee (ERM 29143).

### Pharmacological compounds

Psilocybin (USONA Institute Investigational Drug Supply Program; Lot# AMS0167) was dissolved in saline and administered at a dose of 1.5 mg/kg. Ketanserin tartrate (Tocris Biosciences, CAS 83846-83-7; 1.5mg/kg), MDL100907 (volinanserin; Sigma-Aldrich, CAS 139290-65-6; 0.1mg/kg) and WAY100635 maleate (Tocris Biosciences, CAS 1092679-51-0; 0.5mg/kg) serotonin receptor subtype antagonists were administered 30min before psilocybin (or 0.9% NaCl saline control) treatment and all animals only received one combination of psilocybin/saline and one receptor subtype antagonist. Dose selection was based on the literature ^46, 66–68^. All drugs were administered intraperitoneally at a 1.0ml/kg injection volume using a 26-guage needle.

### Activity-based anorexia (ABA)

The ABA paradigm consists of unlimited access to a running wheel and time-restricted food access. At seven weeks of age, rats were individually housed in transparent living chambers with a removable food basket and a running wheel (Lafayette Instruments, IN, USA). Rats were allowed to habituate to the wheel for seven days to determine baseline running wheel activity (RWA). The following day, psilocybin or saline was administered, wheels were locked for 5h and then reopened. Running activity was recorded by the Scurry Activity Wheel Software (Lafayette Instruments, IN, USA). During the ABA period, food access was restricted to 90min/day at the onset of the dark phase (1100-1230h). Running in the hour before the feeding window (1000-1100h) was considered as food anticipatory activity (FAA). Time-restricted food access persisted for a maximum of 10 days or until rats reached <80% of baseline body weight (ABA susceptibility criterion), at which point they were euthanised with 300mg/kg sodium pentobarbitone (Lethabarb; Virbac, Australia).

### Home-cage operant learning paradigms

Open-source Feeding Experimentation Devices (Version 3), known as “FED3” ^69^, were used for home-cage operant testing, fitted with custom built masks. The task wall consisted of two nose-poke ports situated on either side of a pellet magazine where pellets were delivered with a motorised dispenser. Both operant ports and magazines were fitted with infra-red beams to detect nose-pokes and pellet collection, and were controlled by a commercial microcontroller with data displayed on screen for user feedback. An LED strip underneath the nose-poke ports was used as a light cue. The firmware for FED3 devices were written in the Arduino language, modified from the available Arduino library (https://github.com/KravitzLabDevices/FED3) and flashed in sets of operant training menus (https://github.com/Foldi-Lab/LKM_FED3-tasks).

Following light cycle acclimation, rats were individually housed (26cm W x 21cm H 47.5cm D) with *ad libitum* access to water and standard laboratory chow (Barastoc, Australia) throughout. Rats were habituated to sucrose pellet rewards (20mg, AS5TUT; Test Diet, CA, USA) for two days prior to training. Operant testing was conducted once daily in the home cage for a 3h session between 12:00-15:00 (early dark phase), which began with two days of magazine training on a “free feeding” schedule in which a pellet was dispensed each time one was removed from the magazine. Subsequently, rats were trained to poke for rewards at fixed ratio (FR) schedules (FR1, FR3, FR5) for 2-5 days each until high accuracy (>80% target responding) was achieved. The target side for all experiments was counterbalanced across each cohort to control for any inherent side bias due to in cage FED3 position. Between animal variability in training performance was always balanced between treatment groups and any animals failing to learn the penultimate training step were removed from the experiment before drug administration (see **Supplementary Table 1**).

### Between-session reversal learning task

To test the effects of psilocybin on cognitive flexibility, saline or 5-HTR antagonists (pre-treatment) were administered 30min prior to either saline or psilocybin (treatment), at the completion of the final FR5 training session. The following day (18h post-administration) the reward contingencies of the nose-poke ports were reversed (un-cued), and rats underwent 3 days of testing on the reversed FR5 schedule.

### Fixed and variable ratio schedule training and extinction

To test the effects of psilocybin on suppression of learned FR responding, saline or psilocybin was administered immediately following the final FR5 training session. Over the next 3 days rats underwent extinction testing in which the FED3 was provided as usual except no rewards were delivered regardless of animal activity. To test the effects of psilocybin on training under variable reward schedules, and the long-lasting effects on response suppression, rats were trained to nose-poke at FR1 for 4 days after which saline or psilocybin was administered. The following day rats were trained at variable ratio (VR) schedules of VR5, VR10 and VR20 (two days on each schedule), where the number of target pokes required to deliver a pellet on each trial was randomly selected from 1-5, 6-10 or 11-20, respectively. Subsequently, rats underwent 2 consecutive days of extinction testing, with 24h access to the FED3 device.

### Progressive ratio and re-setting task

To test the effects of psilocybin on motivated (effortful) responding, saline or psilocybin was administered at completion of the final FR5 training session and the next day rats underwent a progressive ratio (PR) reinforcement schedule, where the exponential schedule increased according to the formula (5 * e(0.2*n)-5), where *n* is the trial number, producing response requirements of 1, 2, 4, 6, 9, 12 etc., to the nearest whole number. This was followed by a session at FR5 to reinstate responding and a session during which the PR schedule reset to 1 following any 10min period of FED3 inactivity, called a re-setting progressive ratio (R-PR) task.

### 5-HT receptor subtype abundance

For detection and quantification of 5-HTR subtypes, rats were administered psilocybin or saline and euthanized with sodium pentobarbitone (Lethabarb 150mg/kg; Virbac, AU) at a time course (6, 12 or 24h) post-administration. ABA rats underwent exposure to the model as described above and were administered psilocybin or saline after they had lost at least 15% baseline body weight (15.1-17.6%). Six hours later they were euthanized as above and all rats were transcardially perfused with 200 mL 0.9% saline followed by 200 mL 4% paraformaldehyde in phosphate buffer. Brains were excised and postfixed in 4% paraformaldehyde in phosphate buffer solution overnight at 4°C, followed by submersion in increasing concentrations of 10%, 20% and 30% sucrose in phosphate buffer solution across 3 to 4 days. Brains were then sectioned at 15μm using a cryostat (CM1860; Leica Biosystems) and the medial prefrontal cortex (mPFC) was collected in a 1:4 series. Two mPFC sections per animal, from the same series (spanning from bregma, anteroposterior: +3.2 mm to +2.2 mm), were placed onto SuperFrost Plus slides, and stored at – 20°C until used. The RNAscope ^TM^ Multiplex Fluorescent V2 detection reagent kit (Advanced Cell Diagnostics, USA) was used according to manufacturer’s instructions and included specific *in situ hybridisation* probes complementary to the mRNA of the 5-HT1AR (*Rn-Htr1a*; RDS404801) and 5-HT2AR (*Rn-Htr2a*; ADV424551). Detection of mRNA was achieved using Opal™ fluorophore dyes from the 520 (1:500) and 620 (1:750) reagent packs (Akoya Biosciences, USA). Full protocol details are available in **Supplementary Methods**. Sections were imaged using a widefield microscope (Thunder Imager Live Cell & 3D Assay, Leica Microsystems, Germany) with a PL Fluotar 506007 40x/1.00-0.50-oil Leica objective. The resulting Z-stacks were instantly deconvolved using the integrated Large Volume Computational Clearing deconvolution algorithm (Leica LIGHTNING). The resulting datasets were pre-processed in a custom macro using ImageJ (v1.53t ^70^) and analysed with CellProfiler (v4, ^71^) using a custom pipeline (see **Supplementary Materials**) for quantification of nuclear bodies as well as *Htr1a* and *Htr2a* transcripts. Selected sections were analysed further using Imaris software (v9.9, Oxford Instruments) to establish the anatomical location of identified differences in transcript abundance across the cortical layers ^72^. The DAPI-channel was used as a mask to define individual cells (cell body selection) and the number of *Hrt1a* or *Hrt2a* puncta surrounding DAPI was analysed using the vesicle detection feature.

### Statistical analyses

Statistical analyses were performed using GraphPad Prism 9.5.1 (GraphPad Software, San Diego, CA, USA). Statistical significance was set at *p*<.05, with *p*<.10 considered a trend though not significant. Analyses used were two-tailed unpaired t-test, one-way and two-way analysis of variance (ANOVA) with Bonferroni’s, Dunnett’s or Sidak’s post hoc multiple comparisons, and a mixed-effects model, chosen appropriately considering the type of data, number of groups, and comparisons of interest. Full details of all statistical tests performed (including group composition) can be found in the Statistics Tables in **Supplementary Materials**. For RNAscope analyses each individual animal’s data point represents an average value from 4 (individual regions) or 8 (combined regions) sections.

## Results

### Psilocybin improves body weight maintenance in ABA rats

In order to assess the influence of a single dose of psilocybin on subsequent adaptation to conditions of ABA, psilocybin was administered 24h prior to the onset of timed food restriction, which facilitated improved body weight maintenance throughout ABA exposure (**Fig 1A**) and increased the proportion of animals resistant to the paradigm (**Fig 1B**). Psilocybin-treated rats spent significantly more days above 85% of their baseline body weight (*p*=.0172; **Fig 1C**) and although the reduction in average daily weight loss after psilocybin treatment did not reach statistical significance (*p*=.0638; **Fig 1D**), psilocybin prevented severe weight loss associated with ABA (*p*=.0394; **Fig 1E**). This ability to better maintain body weight under ABA conditions was not driven by marked alterations to overall wheel running (**Fig 1F**), with psilocybin and saline treated animals running similar amounts during both baseline and ABA phases (baseline *p*>.9999, ABA *p*=.3089; **Fig 1G**) and during the food anticipation period (*p*=.2800; **Fig 1H)**. Similarly, food intake increased over successive days of ABA exposure regardless of treatment (**I**) and psilocybin did not change the average amount of food consumed across the ABA period (*p*=.3290; **Fig 1J**). When comparing psilocybin treated rats that were susceptible versus resistant to weight loss, it appeared that psilocybin-induced resistance was not qualitatively distinct from previous work ^58, 73^, but was similarly defined by both reduced food-restriction evoked hyperactivity (**Fig 1K**) that was specific to running during the first ABA (baseline *p*=.7415, ABA *p*<.0001; **Fig 1L**), increased running in anticipation of food (*p*<.0001; **Fig 1M**) and increased food intake across days (**Fig 1N**) and the overall ABA period (*p*<.0001; **Fig 1O**). Notably, the only behavioural feature that preceded improved body weight maintenance after psilocybin treatment was wheel running on the day prior to administration (Baseline Day 7; **Fig 1K**). We also compared only animals treated with psilocybin or saline that were *susceptible* to developing ABA, to understand whether outcomes were altered independent of improved weight maintenance. We found that this subgroup of rats treated with psilocybin were indistinguishable from controls on propensity to engage in starvation induced hyperactivity (**Fig 1P**) that is elicited by ABA (*p*=.2895; **Fig 1Q**), running in anticipation of food (*p*=.5252; **Fig 1R**) or food intake over time (**S**) or on average (*p*=.6368; **Fig 1T**).

**Figure 1.**
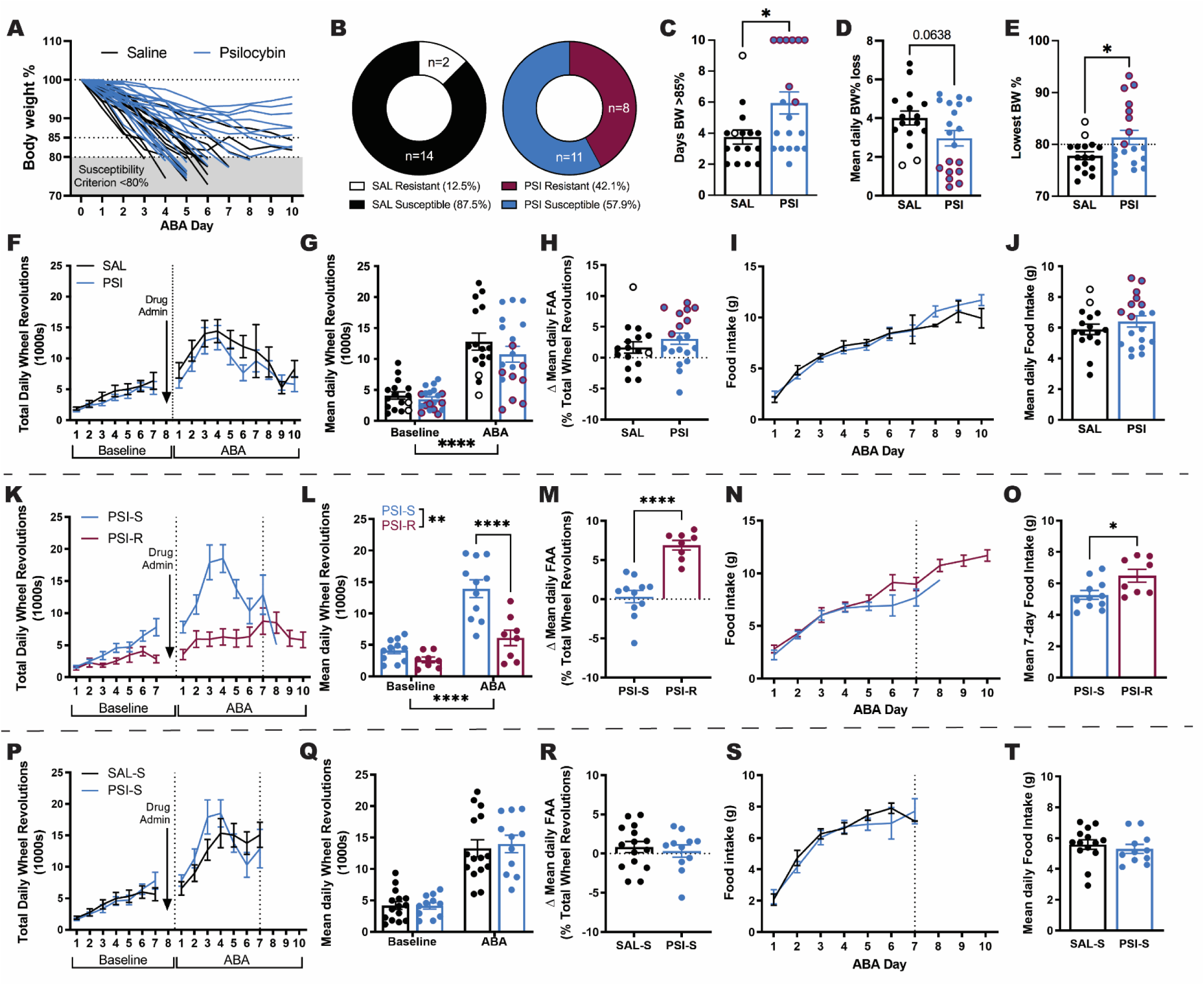
Effects of psilocybin on body weight maintenance in ABA. Weight loss trajectories of individual rats (*n=*16 saline; *n*=19 psilocybin) over the 10-day ABA period (**A**) and proportion resistant to weight loss (**B**). Psilocybin facilitated body weight maintenance over 85% for more days (**C**, *t*(33)=2.508, *p*=.0172), with a trend toward lower body weight % loss per day (**D**, *t*(33)=1.918, *p*=.0638) that resulted in attenuation of severe weight loss (**E**, *t*(33)=2.146, *p*=.0394). Total daily wheel revolutions (**F**) increased as expected in ABA (**G**, ABA Phase *F*(1, 33)=126.5, *p*<.0001) but were similar between groups (Treatment *F*(1, 33)=1.159, *p*=.2985) across both baseline (*p*>.9999) and ABA (*p*=.3089; Interaction *F*(1, 33)=1.033, *p*=.3169) with no difference in the change in proportional running wheel activity in the penultimate hour before food access (**H**, *t*(33)=.1.098, *p*=.2800). Ninety-minute food intake (**I**) increased similarly across the ABA phase with no difference in mean daily intake (**J**, *t*(33)=0.9908, *p*=.3290). Comparison of only <u>psilocybin treated</u> rats that were susceptible (PSI-S) versus resistant (PSI-R) to ABA highlights the characteristic starvation-induced hyperactivity displayed by PSI-S (**K**) during the first 7 days of exposure to ABA conditions (**L**, ABA Phase *F*(1, 17)=36.48, *p*<.0001; ABA Outcome *F*(1, 17)=19.01, *p*<.0001; Interaction *F*(1, 17)=17.09, *p*=.0047; ABA PSI-S>PSI-R *p*<.0001), in contrast to the selective increase of running in anticipation of food access displayed by PSI-R (**M**, *t*(17)=6.203, *p*<.0001), accompanied by diverging food intake trajectories (**N**) with greater mean 7-day intake by PSI-R (**O**, *t*(17)=2.577, *p*=.0196). Comparison of <u>ABA susceptible</u> rats that received psilocybin (PSI-S) or saline (SAL-S) revealed no differences in the development of starvation-induced hyperactivity (**P**) following the onset of ABA conditions (**Q**; ABA Phase *F*(1, 24)=126.5, *p*<.0001; Treatment *F*(1, 24)=1.159, *p*=.2895; Interaction *F*(1, 24)=1.033, *p*=.3269), selective running in anticipation of food (**R**; *t*(24)=0.6452, *p*=05252) or food intake over time (**S**) or on average (**T**; *t*(24)=0.4783, *p*=.6368). Grouped data show mean ± SEM, with individual data points on bar graphs. \**p*<.05, \*\**p*<.01, \*\*\**p*<.001, \*\*\*\**p*<.0001. SAL saline; PSI psilocybin; BW body weight; ABA activity-based anorexia; FAA food anticipatory activity; PSI-S psilocybin treated ABA susceptible; PSI-R psilocybin treated ABA resistant. For full statistical analysis details see Figure 1 **Statistics Table**.

### Psilocybin enhances flexible behaviour in a reversal learning task

Considering that the ability to maintain body weight during exposure to ABA in rats has been previously linked to improved cognitive flexibility on a reversal learning task ^61^ and that exposure to ABA conditions impairs reversal learning ^59^, we hypothesised that the improvements in ABA after psilocybin were associated with improved flexibility in the present study. Psilocybin was administered 18h prior to reversal of reward contingencies (**Fig 2A**), and produced a transient improvement in response accuracy (*p*=.0312; **Fig 2B**), evidenced by a rapid shift in responding towards the reversed port and an increase in the proportion of rats that reached performance criterion (**Fig 2C**). In order to quantify performance, we used a moving window accuracy (80% accurate, within a 100-poke window) to demonstrate that psilocybin treated rats required fewer trials to learn the task (**Fig 2D**). Improved performance after psilocybin was not driven by faster criterion acquisition (*p*=.1474; **Fig 2E**) or altered total (*p*=.1420; **Fig 2F**) or target responses (*p*=.5815; **Fig 2G**), but specifically by reduced responding to the non-target (incorrect) port (*p*=.0260; **Fig 2H**), indicating psilocybin treatment facilitated learning from negative feedback and faster behavioural adaptation, which was also evident in improved reward efficiency (reduced non-target pokes per pellet; see **Supplementary Fig 1F**). While psilocybin did not significantly improve the rate of reward collection (*p*=.0956; **Fig 2I**), it increased engagement with the reversal task evident in reduced latencies to respond (*p*=.0332; **Fig 2J**) and win the first reward (*p*=.0343; **Fig 2K**). To confirm that this improvement was not related to increased effortful responding or response suppression, we tested separate cohorts of rats on progressive ratio (PR), variable ratio (VR) and extinction tasks. Here, we show that psilocybin administration 18h prior to test did not increase the willingness of rats to expend effort to obtain rewards (*p*=.4436; **Fig 2L**), the ability to extinguish a previously learned response (*p*=.5783; **Fig 2M**) or response vigour under uncertain (variable) schedules of reinforcement (*p*=.2013; **Fig 2N**). Moreover, there was no improvement in response suppression 7 days following psilocybin treatment (*p*=.6100; **Fig 2O**). See **Supplementary Fig 1** for full session data, including for animals that did not reach performance criterion on the first reversal session.

**Figure 2.**
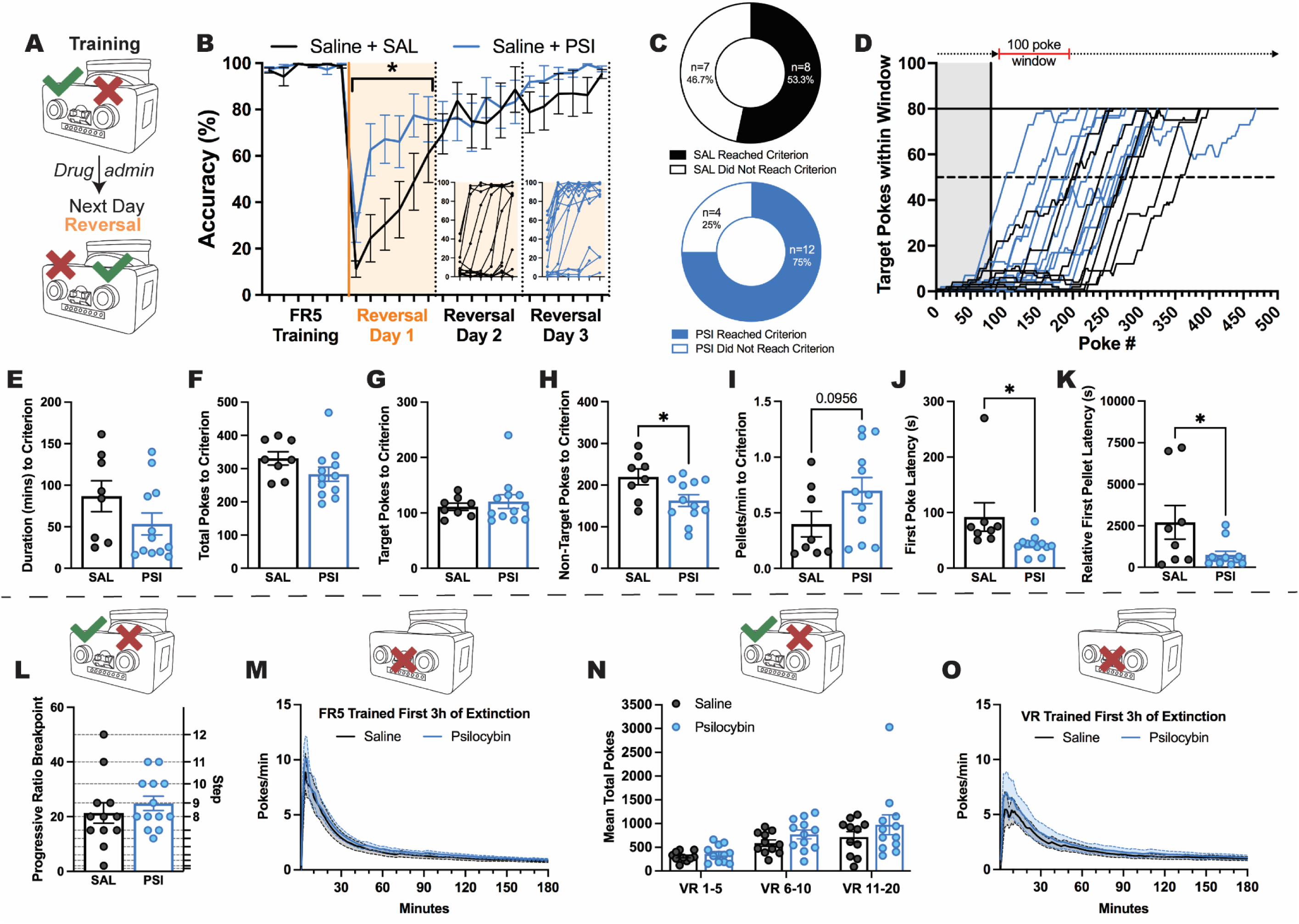
Effects of psilocybin on reversal learning, effortful responding and response suppression. Psilocybin administered after training the day prior to reversal of reward contingencies (**A**) significantly improved accuracy of responding during the initial 3h reversal session (**B**, Treatment *F*(1, 29)=5.128, *p*=.0312; 6 x 30min time bins) and increased the number of rats (**C**) able to reach performance criterion (**D**, 80 target pokes in a 100-poke moving window) on the first day of reversed reward contingencies. While there was no difference in the time (from first poke to poke that achieved criterion; **E**, *t*(18)=1.514, *p*=.1474), total pokes (**F**, *t*(18)=1.536, *p*=.1420) or target pokes (**G**, *t*(18)=0.5614, *p*=.5815) required to reach criterion, psilocybin-treated rats required fewer non-target pokes to reach criterion (**H**, *t*(18)=2.425, *p*=.0260), tended to earn rewards faster (**I**, *t*(18)=1.759, *p*=.0956), and were both faster to first engage with the task (time from device access to first poke; **J**, *t*(18)=2.307, *p*=.0332) and to earn their first reward (time from first poke to earning first pellet; **K**, *t*(18)=2.291, *p*=.0343). Psilocybin treatment had no effect on breakpoint (pokes required to earn final pellet before 10min of inactivity) on a classic progressive ratio task (**L**, *t*(23)=0.7795, *p*=.4436), extinction following fixed ratio training (**M**, Treatment *F*(1, 17)=0.3212, *p*=.5783; Interaction *F*(179, 3043)=0.2623, *p*>.9999), goal directed engagement on increasingly uncertain schedules of reinforcement (**N**, Treatment *F*(1, 21)=1.741, *p*=.2013; Interaction *F*(2, 42)=0.7244, *p*=.4906), or extinction following variable ratio training (**O**, Treatment *F*(1, 21)=0.2681, *p*=.6100; Interaction *F*(179, 3759)=0.3509, *p*>.9999). Grouped data show mean ± SEM, with individual data points on bar graphs. \**p*<.05. SAL saline; PSI psilocybin; FR5 fixed ratio 5; VR variable ratio. For full statistical analysis details see Figure 2 **Statistics Table**.

Because classical PR tasks require the test session to be terminated after a 10 min period of inactivity, and yet psilocybin was shown to increase task engagement in the reversal learning task, we were interested to see if psilocybin also acted to restore responding after periods of inactivity. We tested this in two ways; firstly, by allowing animals access to the operant devices for 3h using a standard PR schedule and secondly, by implementing a variation of the PR task in which after any 10 min period of inactivity the ratio reset to 1 [re-setting PR (R-PR); see **Supplementary Fig 2A-B**]. Breakpoint itself was not different between tasks (all *p*s>.2666; **Supplementary Fig 2C**), however, psilocybin increased task engagement specifically during the R-PR session, when increased engagement is considered economical because the effort required to receive a reward is lower. Moreover, psilocybin-induced task engagement was directed rather than arbitrary, with increases in the number of target but not non-target pokes observed when the ratio reset (PR *p*>.9999, R-PR *p*=.0209; **Supp Supplementary 2D-E**). None of these changes were observed prior to the first re-setting (i.e. first breakpoint; **Supplementary Fig 2I-M**), indicating that experience with the new reward economy was required to elicit increased engagement after psilocybin.

### 5-HT1AR and 5-HT2AR subtype mechanisms differentially drive psilocybin-induced flexible learning

To determine whether psilocybin improved flexibility on the reversal learning task via actions at 5-HT receptor subtypes relevant to anorexia nervosa, selective antagonists to these receptor subtypes were administered 30 min prior to administration of saline (control) or psilocybin and the following day the reward-paired port was reversed (**Fig 3A&L**). Analysis of parameters that contribute to accuracy across the first day of reversal learning (**Fig 3B&M**) revealed that for control rats, 5-HT2AR antagonism completely abolished reversal learning capability, with 0% of rats administered the 5-HT2AR antagonist (MDL100907) reaching performance criterion, compared to approximately 53% of rats administered the 5-HT1AR antagonist (WAY100635) or saline treatment alone (**Fig 3C**). This impairment was driven by all aspects of learning throughout the reversal session, including reduced accuracy (*p*=.0221, **Fig 3D**), rewards obtained (*p*=.0324, **Fig 3E**), target pokes (*p*=.0292, **Fig 3F**) and non-target pokes (*p*=.0048, **Fig 3G**). Importantly, 5-HT2AR antagonism did not cause an impairment in task initiation, since the latency to respond was equivalent across groups (*p*=.4572, **Fig 3H**), although it increased the latency to make a target poke (*p*=.0502, **Fig 3I**), suggesting MDL100907 administration prevented control rats from adapting to the new reward rules. Moreover, the impairment elicited by 5-HT2AR antagonism in control rats was not due to reduced willingness to engage in the task, considering there were no significant changes in the latency to receive the first reward (*p*=.1507, **Fig 3J**) or session duration (*p*=.6048, **Fig 3K**), however, it should be noted that only three MDL100907 treated control rats ever earned rewards. Conversely, 5-HT1A antagonism did not significantly alter most performance measures throughout the test session (all *p*s>.5609; **Fig 3 D-F, H, J, K**) but specifically reduced the number of non-target pokes performed (*p*=.0041, **Fig 3F**) and increased the latency to first target poke (*p*=.0244, **Fig 3I**), suggesting administration of WAY100635 allowed rats to learn to the same degree as saline controls with less negative feedback and despite being slower to respond at the initial reversal of reward contingencies.

**Figure 3.**
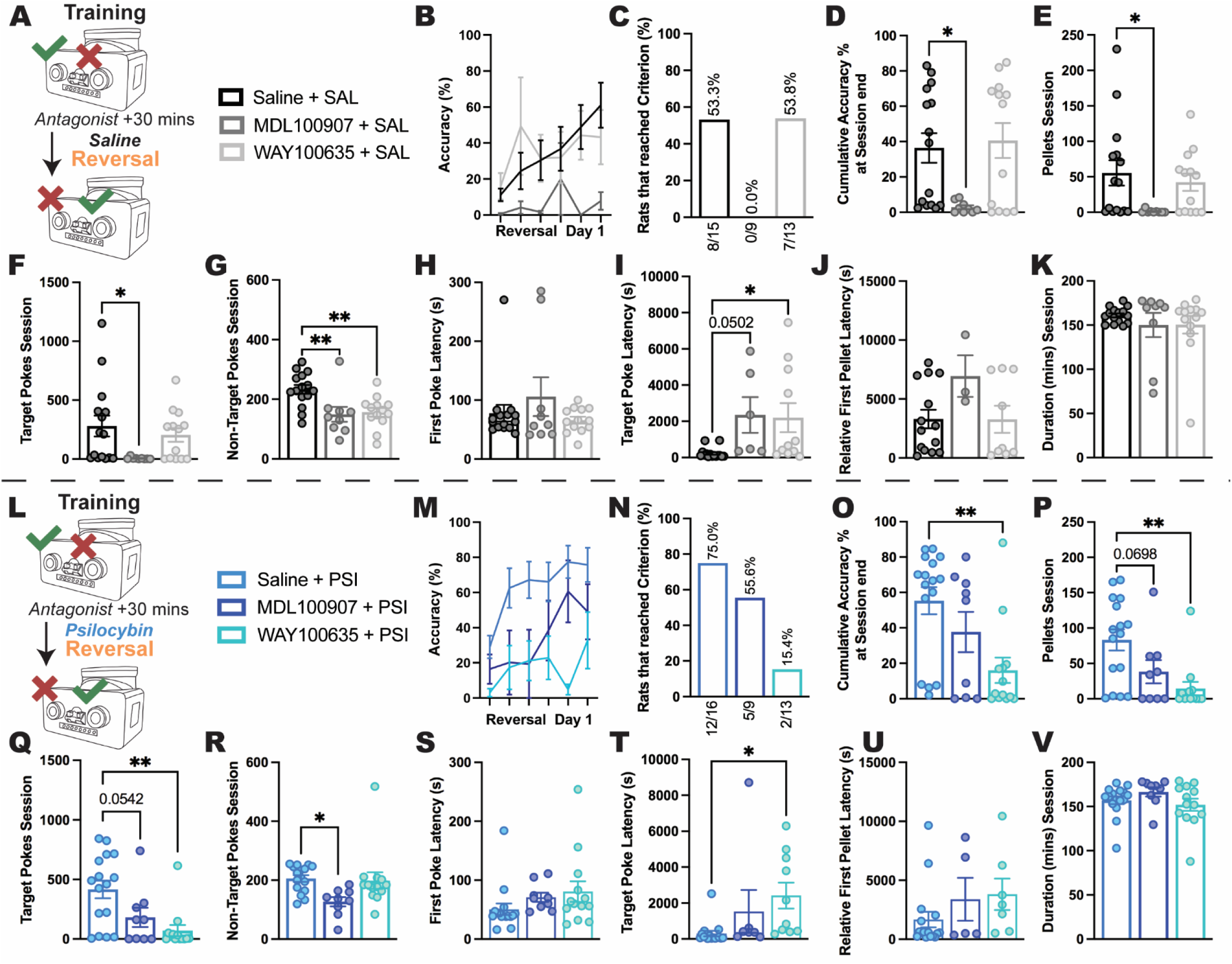
Effects of 5-HT2A and 5-HT1A antagonism on reversal learning in control and psilocybin-treated rats. Following the final training session pre-treatment with either saline (vehicle control), the 5-HT2AR antagonist MDL100907, or the 5-HT1AR antagonist WAY100635, was followed 30min later by treatment with either saline (**A**) or psilocybin (**L**) before reversal of reward contingencies the following day, with first reversal day performance accuracy highlighted (**B and M, respectively).** While 53/5% (8/15) of Saline+SAL rats reached reversal day 1 criterion, 0% (0/9) of MDL+SAL treated rats did so (**C**), showing global impairment compared to Saline+SAL across nearly all outcome measures, achieving significantly lower session accuracy (**D**, SAL+>MDL+ *p*=.0221), earning fewer pellets (**E**, SAL+>MDL+ *p*=.0324), and making fewer target (**F**, SAL+>MDL+ *p*=.0292) and non-target (**G**, SAL+>MDL+ *p*=.0048) pokes in the session. While there was no delay in task engagement (time from device access to first poke; **H**, SAL+ vs MDL+ *p*=.4572), target poke latency was increased (**I**, SAL+<MDL+ *p*=.0502) in the 6/9 rats that made a target poke, and only 3/9 rats earned a single pellet (i.e. made at least 5 target pokes; **J**, time from first poke to earning first pellet, SAL+ vs MDL+ *p*=.1507), even though task engagement duration did not differ (time from first to final poke; **K**, SAL+ vs MDL+ *p*=.6048). Conversely, WAY+SAL resulted in 53.8% (7/13) of rats reaching criterion, nearly identical to Saline+SAL, with these groups being similar across most measures except WAY+SAL having fewer non-target pokes (**G**, SAL+>WAY+ *p*=.0041) despite an elongated target poke latency (**I**, SAL+<WAY+ *p*=.0244). With 75% (12/16) of Saline+PSI rats reaching reversal day 1 criterion, MDL+PSI treatment produced a moderate decrease to 55.6% (5/9) reaching criterion (**N**), although only a non-significant decrease in accuracy (**O**, SAL+ vs MDL+ *p*=.2837), while there was a trend toward fewer pellets (**P**, SAL+>MDL+ *p*=.0698) and target pokes (**Q**, SAL+>MDL+ *p*=.0542), and a significant reduction in non-target pokes (**R**, SAL+>MDL+ *p*=.0210) across the session, with no differences in any latency measures (**S-U**, all SAL+ vs MDL+ *p*s>.2991) or session duration (**V**, SAL+ vs MDL+ *p*=.4345). In contrast, WAY+PSI produced severe impairment with only 15.4% (2/13) of rats reaching criterion, with significantly reduced session accuracy (**O**, SAL+>WAY+ *p*=.0024), pellets earned (**P**, SAL+>WAY+ *p*=.0015), and target pokes (**Q**, SAL+>WAY+ *p*=.0013), and delayed target poke latency (**T**, SAL+<WAY+ *p*=.0212, with only 10/13 rats achieving a target poke) compared to Saline+PSI, whilst there were no differences for non-target pokes (**R**, SAL+ vs WAY+ *p*=.9497), first poke latency (**S**, SAL+ vs WAY+ *p*=.1636), relative first pellet latency (**U**, SAL+ vs WAY+ *p*=.2588, although only 7/13 earned a pellet), nor session duration (**V**, SAL+ vs WAY+ *p*=.7309). Bar graphs show mean ± SEM with individual data points. \**p*<.05, \*\**p*<.01. SAL saline; PSI psilocybin; SAL+ saline pre-treatment; MDL+ MDL100907 pre-treatment; WAY+ WAY100635 pre-treatment. For main ANOVA results and full statistical analysis details see Figure 3 **Statistics Table**.

When combined with psilocybin treatment (**Fig 3L**), antagonism of 5-HT1A and 5-HT2A receptors resulted in an opposing pattern of results, with a substantial reduction in 5-HT1AR antagonist (WAY100635) treated animals able to learn the task to criterion (15.4%; **Fig 3N**), compared to 55.6% 5-HT2AR antagonist (MDL100907) treated and 75% treated with psilocybin alone. This impairment in reversal learning was demonstrated in reduced accuracy (*p*=.0024, **Fig 3O**), rewards obtained (*p*=.0015, **Fig 3P**) and target pokes performed (*p*=.0013, **Fig 3Q**), however, 5-HT1A antagonism prior to psilocybin treatment did not alter suppression of responding to the previously rewarded (non-target) side (*p*=.9497, **Fig 3R**) or willingness to initiate a session (*p*=.1636, **Fig 3S**), although it did increase the latency to poke on the reversed port (*p*=.0212, **Fig 3T**). Compared to psilocybin treatment alone, selective 5-HT2A antagonism reduced the number of non-target pokes (*p*=.0210, **Fig 3R**) but did not significantly alter any other performance measures (all *p*s>.0542; **Fig3 O-Q, S-V**).

### Psilocybin rescues learning impairments induced by 5-HT2AR antagonism potentially via preferential actions at 5-HT1AR

This differential impact of 5-HT2A antagonism is highlighted when comparing performance between saline and psilocybin treated animals that all received MDL100907, a large number of which did not reach performance criterion (**Fig 4A**). Whereas selective 5-HT2A antagonism alone (with saline) impaired performance across the board, co-administration of psilocybin rescued impairments in accuracy (*p*=.0079, **Fig 4B**), rewards earned (*p*=.0385, **Fig 4C**) and target responses (*p*=.0436, **Fig 4D**), potentially via preferential actions at the 5-HT1AR. 5-HT2AR antagonism with MDL100907 administration did not cause differential effects on non-target pokes (*p*=.4578, **Fig 4E**), or the latencies to first poke (*p*=.3163, **Fig 4F**), first target poke (*p*=.6202, **Fig 4G**) or first reward won (*p*=.2428, **Fig 4H**) in psilocybin or saline treated rats, nor was the duration engaged in a session (*p*=.2741, **Fig 4I**) different for psilocybin or saline treated animals administered MDL100907. While 5-HT1A antagonism combined with psilocybin substantially reduced the number of rats able to reach performance criterion (**Fig 4J**), and induced a trend toward reduced accuracy (*p*=.0556, **Fig 4K**), instead of a performance impairment *per se* what seems to be the case is that co-administration of WAY100635 negated psilocybin-induced improvements in reversal learning, with no significant differences in learning measures or response profiles observed between saline and psilocybin treated animals that were administered the 5-HT1AR antagonist (WAY100635) (all *p*s>.0834; **Fig 4L-R)**. Further supporting a role of 5-HT1A antagonism negating the improvement elicited by psilocybin rather than *impairing* performance is the finding that WAY100635 alone facilitated reversal learning compared to saline alone, by reducing non-target responding (**Supplementary Fig 3F**). Intriguingly, co-administration of the mixed antagonist ketanserin impaired performance in saline and psilocybin treated animals to a similar extent, with the notable exception of reducing the session duration for those rats administered saline but not psilocybin (see **Supplementary Fig 4**).

**Figure 4.**
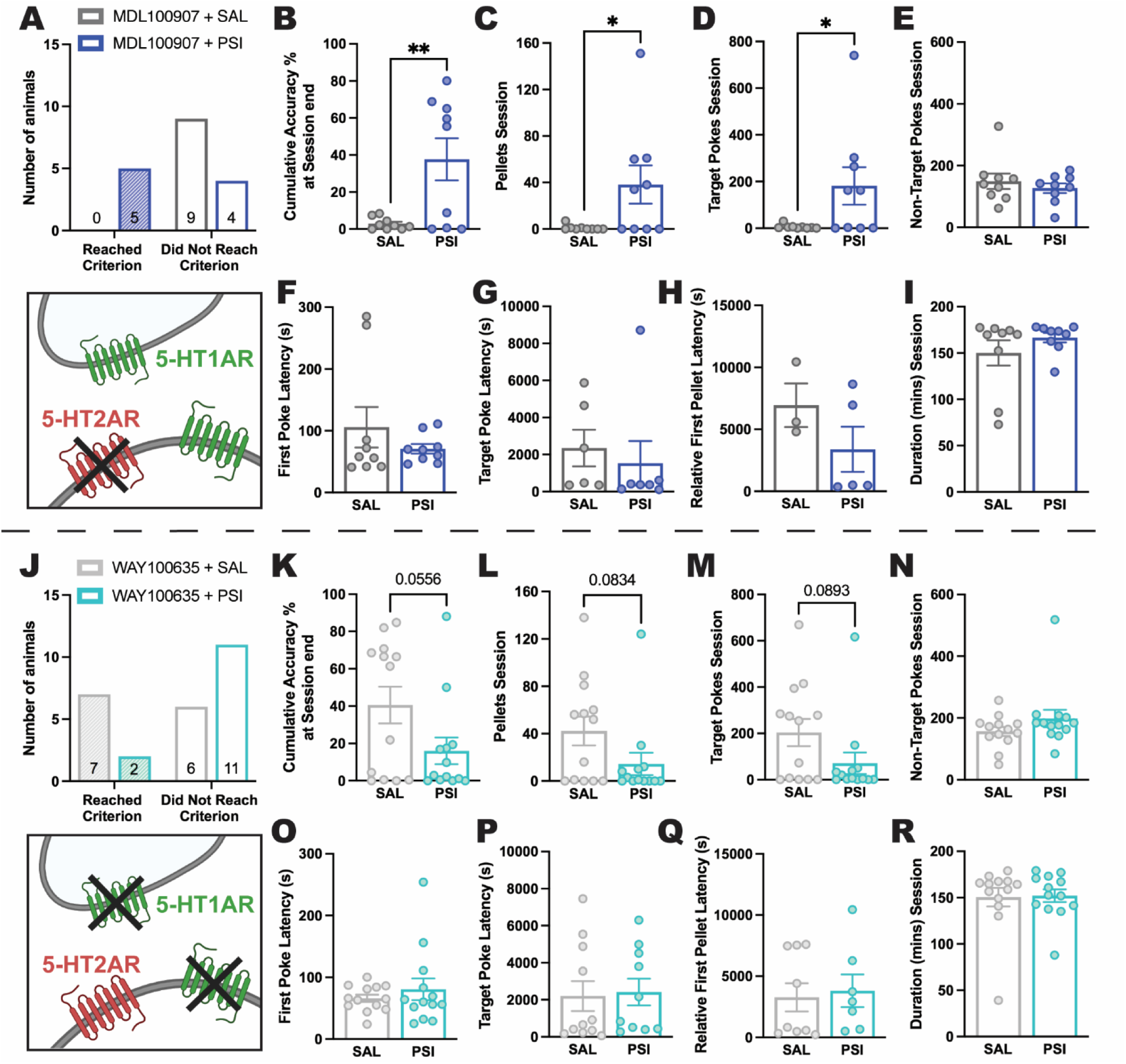
Effects of 5-HT2AR and 5-HT1AR antagonism on psilocybin-induced improvements in reversal learning. Reversal learning following 5-HT2AR antagonism via pre-treatment with MDL100907 (**A**) was completely impaired in saline treated animals (0/9 [0%] reached reversal day 1 criterion) whereas psilocybin treatment prevented this impairment (5/9 [55.6%] reached criterion). Psilocybin treatment following MDL-mediated 5-HT2AR antagonism resulted in significantly greater session accuracy (**B**, *t*(16)=3.034, *p*=.0079), pellets earned (**C**, *t*(16)=2.255, *p*=0.385), and target pokes made (**D**, *t*(16)=2.191, *p*=.0436) compared to saline treatment, with no differences for non-target pokes (**E**, *t*(16)=0.7609, *p*=.4578), first poke latency (time from device access to first poke; **F**, *t*(16)=1.034, *p*=.3163), target poke latency (**G**, *t*(11)=0.5098, *p*=.6202), relative first pellet latency (time from first poke to earning first pellet; **H**, *t*(6)=1.295, *p*=.2428), or session duration (time from first to final poke; **I**, *t*(16)=1.133, *p*=.2741). The opposite performance pattern was observed following 5-HT1AR antagonism via WAY100635 pre-treatment (**J**), with 7/13 (53.8%) saline treated rats reaching criterion compared with only 2/13 (15.4%) psilocybin treated rats. Although not significant, psilocybin treatment produced a trend toward lower session accuracy (**K**, *t*(24)=2.011, *p*=.0556), fewer pellets earned (**L**, *t*(24)=1.806, *p*=.0834), and target pokes made (**M**, *t*(24)=1.771, *p*=.0893), whilst there was no difference between groups for non-target pokes (**N**, *t*(24)=1.317, *p*=.2001), first poke (**O**, *t*(24)=0.7925, *p*=.4538), target poke (**P**, *t*(19)=0.1996, *p*=.8440), or relative first pellet (**Q**, *t*(14)=0.3062, *p*=.7639) latency, or session duration (**R**, *t*(24)=0.1337, *p*=.8948). Bar graphs show mean ± SEM with individual data points. \**p*<.05, \*\**p*<.01. SAL saline; PSI psilocybin. For full statistical analysis details see Figure 4 **Statistics Table**.

### Psilocybin causes a transient shift in the balance of 5-HT1AR and 5-HT2AR mRNA in the prefrontal cortex

To examine whether a change in the abundance of 5-HTR subtypes in medial prefrontal cortex (mPFC) was elicited by psilocybin, which could explain the differential effects of psilocybin on reversal learning after pharmacological blockade of the 5-HT1A vs 5-HT2A receptor subtypes, we performed RNAscope on cortical sections (**Fig 5A**) collected 6, 12 and 24h after psilocybin treatment.

**Figure 5.**
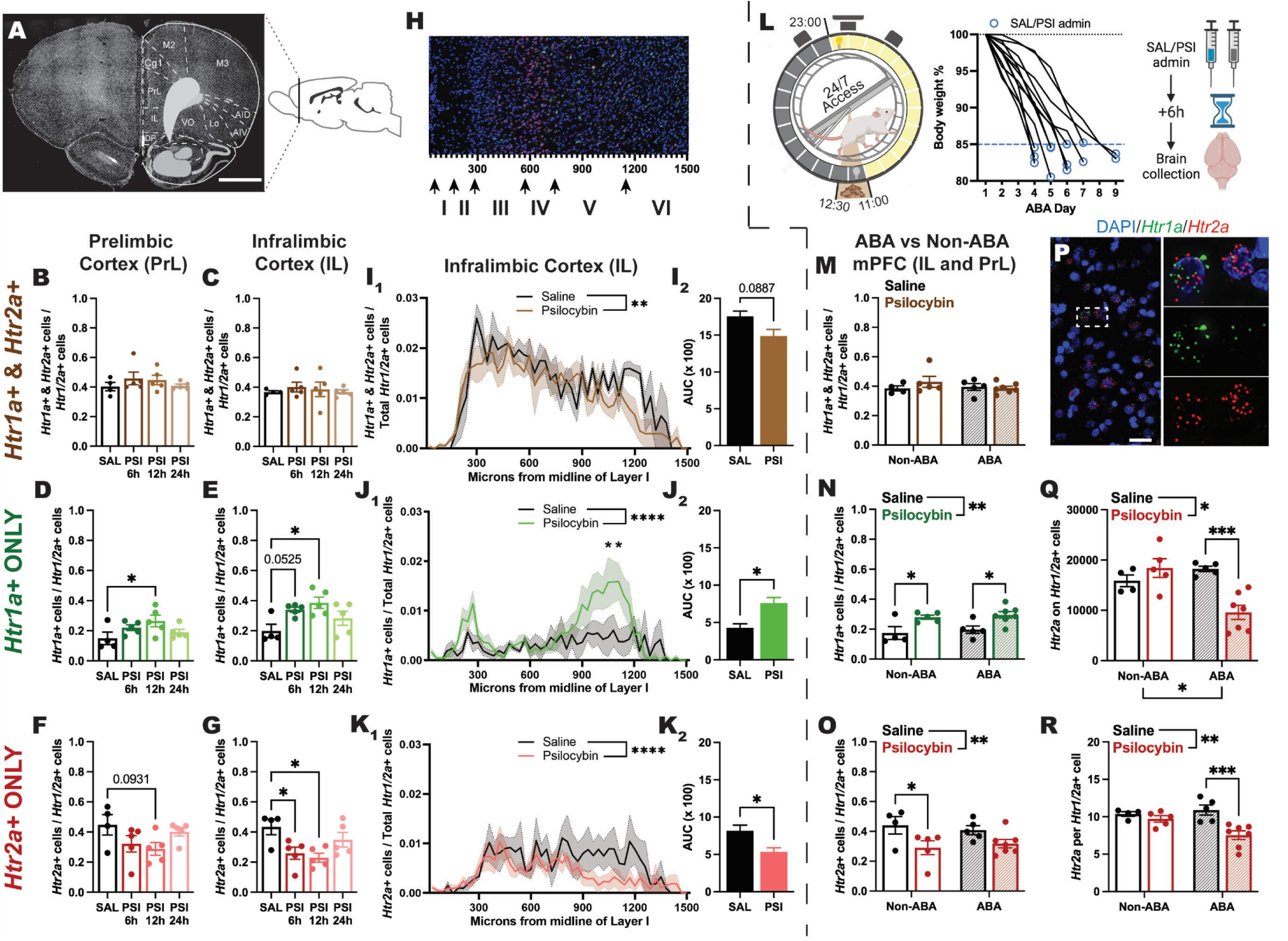
Effects of psilocybin on the expression of *Htr1a* and *Htr2a* transcripts in the mPFC. Coronal section with brain atlas overlay (AP+3.2mm from bregma; **A**) depicting regions of interest (PrL and IL). The proportion of *Htr1/2a*+ cells that were <u>double</u> labelled with *Htr1a* <u>and</u> *Htr2a* was not changed by psilocybin treatment in either the PrL (**B**, *F*(3, 15)=0.7801, *p*=.5233) or IL (**C**, *F*(3, 15)=0.2449, *p*=.8637). The proportion of *Htr1/2a*+ cells that were <u>exclusively</u> *Htr1a* labelled was <u>increased</u> following psilocybin administration in both the PrL (**D**, *F*(3, 15)=2.443, *p*=.1043, SAL<PSI12h *p*=.0500) and the IL (**E**, *F*(3, 15)=4.277, *p*=.0227, SAL<PSI6h *p*=.0525, SAL<PSI12h *p*=.0103), whilst those <u>exclusively</u> *Htr2a* labelled <u>decreased</u> following psilocybin treatment at a trend level in PrL (**F**, *F*(3, 15)=2.192, *p*=.1314, SAL>PSI12h *p*=.0931) and significantly in IL (**G**, *F*(3, 15)=4.426, *p*=.0203, SAL>PSI6h *p*=.0335, SAL>PSI12h *p*=.0129). The spatial distribution of IL *Htr1/2a*+ cells along the midline from Layer I (**H**) was significantly different for each uniquely labelled cell population (**I_1_** *F*(49, 300)=10.34, *p*<.0001; **J_1_** *F*(49, 300)=3.549, *p*<.0001; **K_1_** *F*(49, 300)=3.436, *p*<.0001). In each case psilocybin treatment also had a significant effect, producing a significantly reduced overall proportion of <u>double</u> labelled cells (**I_1_**, *F*(1, 300)=9.214, *p*=.0026) and <u>exclusively</u> *Htr2a* labelled cells (**K_1_**, *F*(1, 300)=19.16, *p*<.0001), but a significantly increased overall proportion of <u>exclusively</u> *Htr1a* labelled cells (**J_1_**, *F*(1, 300)=22.38, *p*<.0001) accompanied by a significant Distance by Treatment interaction (**J_1_**, *F*(49, 300)=1.459, *p*=.0313). AUC was decreased at a trend level for <u>double</u> labelled cells (**I_2_**, *t*(6)=2.030, *p*=.0887), significantly increased for <u>exclusively</u> *Htr1a* labelled cells (**J_2_**, *t*(6)=3.102, *p*=.0211) and significantly <u>reduced</u> for <u>exclusively</u> *Htr2a* labelled cells (**K_2_**, *t*(6)=3.097, *p*=.0212). A separate cohort of animals underwent ABA induction, were administered either saline or psilocybin when they reached <85% baseline body weight, and culled ∼6h later (when bodyweight had dropped to close to ∼80% in most cases; **L**). The proportion of mPFC *Htr1/2a*+ cells that expressed <u>both</u> *Htr1a* and *Htr2a* was not effected by psilocybin administration nor ABA exposure (**M**, Treatment *F*(1, 17)=0.5663, *p*=.4620; ABA Exposure *F*(1, 17)=0.4685, *p*=.5029; Interaction *F*(1, 17)=1.208, *p*=.2871), whereas psilocybin significantly <u>increased</u> or significantly <u>decreased</u> the proportion of <u>exclusively</u> *Htr1a* labelled (**N**, Treatment *F*(1, 17)=15.50, *p*=.0011; Non-ABA SAL<PSI *p*=.0298, ABA SAL<PSI *p*=.0206) or *Htr2a* labelled (**O**, Treatment *F*(1, 17)=9.038, *p*=.0079; Non-ABA SAL>PSI *p*=.0463) cells, respectively, in a generally consistent and ABA independent manner (all ABA exposure and interaction *p*s>.4607). *Htr1a* (green) and *Htr2a* (red) expression on distinct cell populations in the mPFC (**P**) identified through DAPI (blue). The absolute number of *Htr2a* transcripts associated with mPFC *Htr1/2a*+ cells (**Q**) was significantly altered by psilocybin (*F*(1, 17)=4.587, *p*=.0470), ABA exposure (*F*(1, 17)=5.098, *p*=.0374), and their interaction (*F*(1, 17)=15.26, *p*=.0011), such that psilocybin treatment significantly reduced *Htr2a* copy number specifically in the ABA brain (ABA SAL>PSI *p*=.0005). This pattern was mostly replicated by the number of *Htr2a* copies per mPFC *Htr1/2a*+ cell (**R**, Treatment *F*(1, 17)=12.12, *p*=.0029; ABA Exposure *F*(1, 17)=2.048, *p*=.1705; Interaction *F*(1, 17)=5.606, *p*=.0300), with a significant reduction in the density of *Htr2a* specifically following psilocybin treatment after ABA induction (ABA SAL>PSI *p*=.0007). Grouped data show mean ± SEM, with individual data points on bar graphs (except AUC). Values are the average of 4 (PrL and IL) or 8 (mPFC) sections per animal. \**p*<.05, \*\**p*<.01, \*\*\**p*<.001, \*\*\*\**p*<.0001. AP anterior-posterior; SAL saline; PSI psilocybin; PrL prelimbic cortex; IL infralimbic cortex; AUC area under the curve; mPFC medial prefrontal cortex (PrL and IL combined); *Htr1/2a*+ cells expressing *Htr1a* and/or *Htr2a*; ABA activity-based anorexia. Scale bars for **A**) 2mm and **P**) 30µm. For full statistical analysis details see Figure 5 **Statistics Table**.

Psilocybin had no effects on the proportion of cells positive for both *Htr1a* and *Htr2a* transcripts in either the prelimbic (*p*=.5233, **Fig 5B**) or infralimbic (*p*=.8637, **Fig 5C**) subregions of the mPFC, but significantly increased the proportion of cells exclusively positive for *Htr1a* in both subregions (prelimbic; *p*=.0500, **Fig 5D**, infralimbic; *p*=.0103, **Fig 5E**) at 12h post-administration. This was matched with complementary reductions elicited by psilocybin in the proportion of cells exclusively positive for *Htr2a* in both subregions, although this did not reach statistical significance for the prelimbic cortex (*p*=.0931, **Fig 5F**) and was evident at both 6h and 12h timepoints for infralimbic cortex (6h; *p*=.0335, 12h; *p*=.0129, **Fig 5G**). We further examined the anatomical localisation of *Htr1a* and *Htr2a* positive cells across the layers in infralimbic cortex (**Fig 5H**) at 12h post-administration to show an overall reduction in double positive cells after psilocybin treatment (*p*=.0026, **Fig 5I_1_**) that did not reach significance when analysed as an area under the curve (AUC; *p*=.0887, **Fig 5I_2_**). What was clear, however, was that the respective increase and decrease in the proportion of cells exclusively positive for *Htr1a* or *Htr2a* 12h after psilocybin treatment was specifically localised to cortical Layer V (*Htr1a p*<.0001, **Fig 5J_1_,** *p*=.0211, **Fig 5J_2_**; *Htr2a p*<.0001, **Fig 5K_1_** *p*=.0212, **Fig 5K_2_**), which corresponded to 900-1200 µm distance from the midline of Layer I (see **Fig 5H**).

To determine whether psilocybin had similar effects on *Htr1a* and *Htr2a* expression in the context of weight loss and feeding pathology relevant to anorexia nervosa, we compared transcripts from the saline and 6h psilocybin treated rats (non-ABA) to rats that had exhibited substantial weight loss after exposure to ABA conditions (and brains collected 6h after psilocybin administration) (**Fig 5L**). Main effects of psilocybin identified in non-ABA rats were recapitulated for ABA rats, with no changes in the number of double labelled cells (*p*=.4620, **Fig 5M**) but complementary increases in *Htr1a* (*p*=.0011, **Fig 5N**) and decreases in *Htr2a* (*p*=.0079, **Fig 5O**) positive cells, indicating similar consequences of psilocybin treatment occurred in the ABA brain. Multiple comparisons revealed that the increase in the number of *Htr1a* positive cells elicited by psilocybin was stronger in ABA rats than non-ABA rats (non-ABA; *p*=.0298, ABA; *p*=.0206, **Fig 5N**), while the decrease in *Htr2a* positive cells was weaker (non-ABA; *p*=.0463, ABA; *p*=.2102, **Fig 5O**). However, additional changes in the abundance of *Htr2a* transcripts (**Fig 5P**) were observed following weight loss under ABA conditions, whereby psilocybin elicited a substantial reduction in the overall number of *Htr2a* transcripts (*p*=.0005, **Fig 5Q**) and in the number of *Htr2a* transcripts per cell (*p*=.0007, **Fig 5R**) in the mPFC of ABA rats, that was not evident in non-ABA rats. Further analyses of changes in *Htr1a* and *Htr2a* expression over the 24h time course and effects of exposure to ABA are provided in **Supplementary Fig 5**.

## Discussion

Clinical trials evaluating the safety and efficacy of psilocybin in people with AN have been ongoing since 2019, with the first pilot study recently reporting that it improves eating disorder symptoms in some individuals ^74^. Psilocybin may have transdiagnostic efficacy^18, 31, 40^ through several mechanisms relevant to the pathology of AN, including actions on the serotonergic system ^39^ and cognitive flexibility ^22^. However, the details of how such mechanisms are altered by psilocybin in the context of AN remains unknown. Here, we show that psilocybin improves body weight maintenance in the ABA rat model and enhances cognitive flexibility in a reversal learning task by both reducing perseverative responding and promoting task engagement when reward contingencies are initially reversed. That psilocybin did not elicit changes in motivated responding (PR) or response suppression (extinction) following the same training and drug administration protocol suggests a selective improvement in adaptive cognition in the face of changing rules.

Further, we demonstrate that psilocybin-induced improvements in reversal learning performance were not dependent on binding to the 5-HT2AR, because co-administration of the selective 5-HT2AR antagonist (MDL100907) did not significantly alter performance measures. Instead, the action of psilocybin at the 5-HT1AR was required for improved cognitive flexibility, whereby improvements in reversal accuracy and engagement were abolished when psilocybin was co-administered with the selective 5-HT1AR antagonist (WAY100635). This finding is complicated by the relatively fast-acting effects of psilocybin observed on *Hrt2a and Hrt1a* transcription in the mPFC, which indicates that psilocybin rapidly and transiently alters the balance of the cellular machinery required to support receptor binding in this region associated with cognitive flexibility ^49^. In such a way, the differential effects of 5-HT2A and 5-HT1A antagonism on reversal learning after psilocybin may reflect functional interactions between these two receptor subtypes that depend on serotonin availability during the post-acute (∼24h) administration period ^75^. These outcomes also call into question reports of the necessity of 5-HT2A binding for “therapeutic” outcomes of psilocybin in animal models, particularly those that use the non-selective antagonist, ketanserin. Not only does ketanserin bind multiple serotoninergic and non-serotonergic receptors but it also only blocks ∼30% of 5-HT2AR in the rat cortex ^76^. It is plausible, therefore, that partial blockade with ketanserin shifts the binding of psilocybin to other 5-HTR subtypes, including 5-HT1A, which may explain the acute improvement in reversal learning after ketanserin alone previously reported in rats ^38^.

The finding that psilocybin administration specifically prevented severe weight loss in ABA rats is critical in light of the evidence that lower body mass increases the risk for fatal outcomes in AN ^77, 78^. That psilocybin treatment did not have overall effects on feeding or exercise independently is unsurprising considering that psilocybin does not alter feeding or energy balance in mouse models of obesity ^79^ and supports the proposal that the therapeutic effects of psilocybin for anorexia nervosa are more likely driven by adaptive cognition than through metabolic alterations ^53^. In line with this, resistance to weight loss after psilocybin was associated with all aspects of behavioural adaptation to ABA conditions (i.e. increased food intake, reduced compulsive running and increased motivated running), which we have previously shown to be linked with improved cognitive flexibility in ABA rats ^59, 61^. While we only observed trend level reductions in overall body weight loss after psilocybin administration, the treatment group is clearly comprised of two distinct subgroups – those that respond to psilocybin with improved weight outcomes and those that are indistinguishable from controls. This divergence in response profiles exists in multiple clinical populations, where between 40-80% of individuals report therapeutic benefits of psilocybin assisted psychotherapy at follow-up, dependent on trial parameters ^16, 74, 80^. Response variation was also seen in the pilot study of psilocybin in people with AN, with clinically significant improvements seen in 4/10 participants ^74^. The effects of psilocybin on ABA and cognitive flexibility were not assayed in the same subjects in the present study, due to confounds associated with using food rewards in a model that is typified by disturbed feeding behaviour and dysregulated reward processing ^81^. However, considering the specific effects of psilocybin on perseverative behaviour during reward reversal observed, perhaps those individuals (humans or rats) who respond to psilocybin with positive body weight outcomes represent a subgroup whose profile is typified by rigid patterns of thought and behaviour. This information could guide the clinical application of psilocybin to those individuals demonstrating high rigidity. In a similar vein, the observation that rats that responded to psilocybin with improved weight outcomes demonstrated lower levels of running during the baseline phase (i.e. prior treatment and the onset of ABA conditions; see **Supplementary Fig 6**) points to the intriguing potential that psilocybin may be more efficacious in animals (and possibly people) that already have a lower propensity to engage in excessive exercise.

The translational relevance of administering psilocybin prior to ABA exposure in this study, instead of after the establishment of anorectic phenotypes (as is the case for the clinical situation) requires some elaboration. Because of the ethical requirement to remove animals from the ABA paradigm when they reach the weight loss criterion that deems them “susceptible”, it is not possible to intervene at this point to attempt to improve outcomes. We attempted to delay administration after a specific duration of ABA exposure (2 days) or after a specific amount of weight loss (15%) in separate cohorts of rats, however, this intervention produced worse outcomes for both saline and psilocybin treated rats, likely due to the additional stress associated with handling and injection at this critical point of the ABA paradigm (see **Supplementary Methods & Supplementary Fig 7**). It is also important to recognise the short generation time for the ABA phenotype compared to the often long and protracted pathogenesis of AN ^82^ that may underscore differences in the timing and nature of impairments in cognitive flexibility between human and rodent. Whereas cognitive inflexibility may exist prior to onset of AN symptoms and contribute to the development of the condition, it does not predict susceptibility to ABA but develops coincident with weight loss in rats ^59^. Thus, the effects of prior administration of psilocybin on weight maintenance has relevance for the specific type of inflexibility that develops in the context of eating pathology in the rat model. In both cases (human and rat) further research is required to understand how psilocybin might elicit meaningful changes in body weight maintenance over the long term, and to determine whether the same mechanisms underpin the effects of psilocybin on body weight maintenance and cognitive flexibility ^83^.

The specific focus in the present study on the involvement of 5-HT2A and 5-HT1A receptor subtypes was based in the evidence from imaging studies that AN is associated with decreased 5-HT2A and increased 5-HT1A binding in cortical regions ^23, 24^. The finding that psilocybin has the same main effects on the number of cortical cells exclusively positive for the *Htr1a* and *Htr2a* transcripts in animals that had been exposed to ABA conditions is important for the clinical application of psilocybin for AN, and suggests that at least some of the neurobiological effects are unchanged by the development of AN-relevant symptoms. Notably, psilocybin treatment in ABA rats was associated with an augmented increase in the number of cells exclusively positive for *Htr1a* transcripts and an additional reduction in the abundance *Htr2a* transcripts (i.e. number of transcripts per cell) that was not seen after psilocybin treatment in rats that were naïve to ABA. This suggests that in the context of AN-associated symptoms, the actions of psilocybin on cellular activity in the mPFC is more inhibitory in nature, which could indeed be therapeutically relevant in light of the evidence that AN is associated with exaggerated cortical activity ^84^. Perhaps then, it is this additional boost of inhibitory tone elicited by psilocybin in ABA rats that allows them to better adapt to the experimental conditions of ABA when psilocybin treatment is administered prior to onset.

The overall influence of psilocybin on the number of mPFC cells that exclusively express *Htr1a* and *Htr2a* transcripts is also relevant for understanding the involvement of activity in this brain region for cognitive inflexibility in ABA rats. If one considers the large majority (60-75%) of mPFC Layer V cells (where the effects of psilocybin were localised) that express these mRNAs are pyramidal (glutamatergic) cells, the net effect of psilocybin during this 12h window would be hyperpolarisation of the mPFC, via both increasing the inhibitory 5-HT1AR and decreasing the excitatory 5-HT2AR machinery. This aligns with our previous work, in which chemogenetic suppression of mPFC projection neurons could both prevent weight loss in ABA and improve flexibility on a reversal learning task ^61^. However, *Htr1a* and *Htr2a* transcripts are also present on at least two classes of GABAergic interneurons in this cortical region, complicating the interpretation of effects of psilocybin on excitatory output ^85^. Moreover, 5-HT1AR are expressed both pre- and post-synaptically, with differential effects of binding on serotonergic transmission ^86, 87^. Finally, the alterations observed at a transcriptional level does not preclude other mechanisms such as protein degradation or changes in receptor cycling ^88, 89^ from being involved in the serotonergic consequences of psilocybin treatment.

With respect to the specific improvement in reversal learning elicited by psilocybin as a mechanism to explain improved body weight maintenance during ABA, it is notable that the reduced perseverative responding when reward contingencies were reversed was also driven by a subpopulation of “responders”. This raises the possibility that individual differences in baseline serotonin signalling may underlie responses to psilocybin treatment, as proposed by the inverted “U-shaped” dose-effect relationships reported for many active compounds and their relation to cognitive function ^90^. If adaptive cognition requires an appropriate balance between 5-HT1A and 5-HT2A receptor function ^49^, our molecular findings suggest that individuals exhibiting elevated 5-HT1AR function (or for that matter reduced 5-HT2AR function) may not respond positively (since further elevation or reduction elicited by psilocybin would push them into the tail ends of the inverted “U”). It is also important to note, in light of the recent observation that psilocybin, administered acutely, did not facilitate flexibility ^38^, that there are important methodological differences that may explain this discordance. Specifically, we examined effects of psilocybin post-acutely, using a single administration paradigm, and the reversal learning task used in the present study relied on action-outcome learning rather than Pavlovian cue-outcome learning. Performance on this task is also dependent on the incentive salience of rewards to elicit appropriate responding, with psilocybin-induced improvements observed in reversal task engagement, leading to faster receipt of the first (unexpected) reward. This demonstrates the potential of psilocybin to alter the explore/exploit trade-off common in reinforcement learning, where the subject has the option of maximizing reward based on its current information (exploitation) or by accumulating more evidence (exploration) ^91^ and may improve the balance between the two for more effective adaptation.

One of the most intriguing issues related to the actions of psilocybin in the brain is the means via which it changes neuronal morphology and function to exert its effects. The canonical pathway through which psilocybin is proposed to promote plasticity (and presumably therefore flexible learning) is through binding to the 5-HT2A receptor, an act that elicits a “glutamate surge” through rapid increases in neuronal excitability ^29, 30^. The dendritic and synaptic changes that occur downstream may or may not be related to this surge of glutamate since psilocybin induced structural plasticity was still observed in the presence of ketanserin ^46^. It is convenient to focus the actions of psilocybin at 5-HT2AR located in the PFC because of their requirement for the subjective (psychedelic) effects ^43^, however, this view discounts the abundant expression of 5-HT2A in other brain regions relevant to learning and memory, including the hippocampus, claustrum and striatum ^92^. The results of the present study suggest that improvements in flexible learning after psilocybin are not mediated by binding to the 5-HT2A receptor, but that selective 5-HT2A antagonism impaired learning in all animals, an effect that was partially restored with co-administration with psilocybin. A possible explanation for these results is that while 5-HT2A receptor function is required for reversal learning, it only partly supports the cognitive enhancing effects of psilocybin. A major challenge is in understanding the role of the 5-HT1A receptor in mediating learning outcomes, especially since firing activity of 5-HT neurons in the dorsal raphe nucleus is controlled by pre-synaptic expression of 5-HT1AR where binding inhibits serotonin release ^92^. We show that 5-HT1A antagonism did not affect the ability to reach performance criteria or obtain reward in controls, but preferentially impaired learning improvements elicited by psilocybin. Taken together, this highlights the 5-HT1AR as an important target mediating the effects of psilocybin on cognitive flexibility ^92^.

The key outcomes of this study underline the fact that animal studies are required for understanding the mechanisms that underlie the therapeutic efficacy of psilocybin because they allow detailed interrogation of behaviour and brain function in the absence of effects of expectancy. It is important to note that female animals were used exclusively in these studies and that psilocybin has been shown to have effects that differ based on biological sex in rats and mice in terms of behaviour ^93^, regional brain reactivity ^94^ and structural neuroplasticity ^46^. Future studies should aim to examine how psilocybin influences serotoninergic tone via other (i.e. non-5-HT2A) known 5-HT binding targets in both male and female animals. In addition, real-time longitudinal monitoring of 5-HT activity (i.e. with fiber photometry) could be employed in combination with pharmacological tools to shed light on the 5-HT2AR drivers of weight loss in ABA rats, by measuring dynamic changes as animals progress through the paradigm. This would allow precise differentiation between the effects of psilocybin on susceptible versus resistant subgroups, information that would be particularly relevant for the clinical application of psilocybin in individuals with AN. Additionally, examination of the interaction between serotonergic and dopaminergic mechanisms that influence the way that inflexible patterns of thought and behaviour relate to food reward, aversion, and avoidance ^95^ should be a focus for future research. That psilocybin has direct actions on the dopamine system in humans ^96^ and rats ^51^ has long been known, but surprisingly paid little attention ^97^, even though the interaction between serotonin receptor binding and dopamine release is well established ^98^. The proposal to study dopaminergic effects of psilocybin is brought into sharper focus by recent evidence of brain-wide activation of the dopamine system by ketamine ^99^ and that dopamine D2 receptor blockade attenuates the psychedelic-induced head-twitch response ^100^. These considerations, in concert with the new data presented here, will provide a better understanding of a mechanistic framework of psilocybin actions in the brain, insight that will provide greater confidence in the potential therapeutic use of psilocybin for conditions such as AN. This is an important and arguably necessary step towards including psilocybin in the armoury of tools to treat mental health disorders.

## Acknowledgements

We acknowledge the USONA Institute Investigational Drug Supply Program for providing the psilocybin used in these studies and A/Prof Alexxai Kravitz, Washington University in St. Louis for input into operant task design and operations. We acknowledge the use of the Monash Microimaging Facility and Biorender for aspects of the figures.

## Funding

This work was supported by the National Health and Medical Research Council (NMHRC) of Australia (Ideas Grant; GNT2011334 – CJF).

## Conflict of interest

CJF sits on the scientific advisory board for Octarine Bio, Copenhagen, Denmark.

## Supplementary materials

Supplementary information is available in a separate document attached to this submission.

## Notes

### Summary of Updates

Additional analyses between animals that were susceptible to weight loss during exposure to ABA treated with psilocybin or saline (Figure 1; lower panel). Additional information in the supplement about the delayed dose protocol for effects of psilocybin on ABA (Supplementary Figure 7).

## References

1. Chesney E, Goodwin GM, Fazel S. Risks of all-cause and suicide mortality in mental disorders: a meta-review. World Psychiatry 2014; 13(2): 153–160.

2. Arcelus J, Mitchell AJ, Wales J, Nielsen S. Mortality rates in patients with anorexia nervosa and other eating disorders. A meta-analysis of 36 studies. Arch Gen Psychiatry 2011; 68(7): 724–731.

3. van Eeden AE, van Hoeken D, Hoek HW. Incidence, prevalence and mortality of anorexia nervosa and bulimia nervosa. Curr Opin Psychiatry 2021; 34(6): 515–524.

4. Ferguson CP, La Via MC, Crossan PJ, Kaye WH. Are serotonin selective reuptake inhibitors effective in underweight anorexia nervosa? Int J Eat Disord 1999; 25(1): 11–17.

5. Miles S, Phillipou A, Sumner P, Nedeljkovic M. Cognitive flexibility and the risk of anorexia nervosa: An investigation using self-report and neurocognitive assessments. J Psychiatr Res 2022; 151: 531–538.

6. Miles S, Gnatt I, Phillipou A, Nedeljkovic M. Cognitive flexibility in acute anorexia nervosa and after recovery: A systematic review. Clin Psychol Rev 2020; 81: 101905.

7. Smith KE, Mason TB, Johnson JS, Lavender JM, Wonderlich SA. A systematic review of reviews of neurocognitive functioning in eating disorders: The state-of-the-literature and future directions. Int J Eat Disord 2018; 51(8): 798–821.

8. Wu M, Brockmeyer T, Hartmann M, Skunde M, Herzog W, Friederich HC. Set-shifting ability across the spectrum of eating disorders and in overweight and obesity: a systematic review and meta-analysis. Psychol Med 2014; 44(16): 3365–3385.

9. Tchanturia K, Anderluh MB, Morris RG, Rabe-Hesketh S, Collier DA, Sanchez P et al. Cognitive flexibility in anorexia nervosa and bulimia nervosa. J Int Neuropsychol Soc 2004; 10(4): 513–520.

10. Tchanturia K, Davies H, Roberts M, Harrison A, Nakazato M, Schmidt U et al. Poor cognitive flexibility in eating disorders: examining the evidence using the Wisconsin Card Sorting Task. PLoS One 2012; 7(1): e28331.

11. Brockmeyer T, Febry H, Leiteritz-Rausch A, Wunsch-Leiteritz W, Leiteritz A, Friederich HC. Cognitive flexibility, central coherence, and quality of life in anorexia nervosa. J Eat Disord 2022; 10(1): 22.

12. Dajani DR, Uddin LQ. Demystifying cognitive flexibility: Implications for clinical and developmental neuroscience. Trends Neurosci 2015; 38(9): 571–578.

13. Uddin LQ. Cognitive and behavioural flexibility: neural mechanisms and clinical considerations. Nat Rev Neurosci 2021; 22(3): 167–179.

14. Carhart-Harris RL, Bolstridge M, Day CMJ, Rucker J, Watts R, Erritzoe DE et al. Psilocybin with psychological support for treatment-resistant depression: six-month follow-up. Psychopharmacology (Berl) 2018; 235(2): 399–408.

15. Carhart-Harris RL, Bolstridge M, Rucker J, Day CM, Erritzoe D, Kaelen M et al. Psilocybin with psychological support for treatment-resistant depression: an open-label feasibility study. Lancet Psychiatry 2016; 3(7): 619–627.

16. Johnson MW, Garcia-Romeu A, Griffiths RR. Long-term follow-up of psilocybin-facilitated smoking cessation. Am J Drug Alcohol Abuse 2017; 43(1): 55–60.

17. Moreno FA, Wiegand CB, Taitano EK, Delgado PL. Safety, tolerability, and efficacy of psilocybin in 9 patients with obsessive-compulsive disorder. J Clin Psychiatry 2006; 67(11): 1735–1740.

18. IsHak WW, Garcia P, Pearl R, Dang J, William C, Totlani J et al. The Impact of Psilocybin on Patients Experiencing Psychiatric Symptoms: A Systematic Review of Randomized Clinical Trials. Innov Clin Neurosci 2023; 20(4-6): 39–48.

19. Vollenweider FX, Preller KH. Psychedelic drugs: neurobiology and potential for treatment of psychiatric disorders. Nat Rev Neurosci 2020; 21(11): 611–624.

20. Psychedelics bind to TrkB to induce neuroplasticity and antidepressant-like effects. Nat Neurosci 2023; 26(6): 926–927.

21. Vargas MV, Dunlap LE, Dong C, Carter SJ, Tombari RJ, Jami SA et al. Psychedelics promote neuroplasticity through the activation of intracellular 5-HT2A receptors. Science 2023; 379(6633): 700–706.

22. Doss MK, Povazan M, Rosenberg MD, Sepeda ND, Davis AK, Finan PH et al. Psilocybin therapy increases cognitive and neural flexibility in patients with major depressive disorder. Transl Psychiatry 2021; 11(1): 574.

23. Audenaert K, Van Laere K, Dumont F, Vervaet M, Goethals I, Slegers G et al. Decreased 5-HT2a receptor binding in patients with anorexia nervosa. J Nucl Med 2003; 44(2): 163–169.

24. Bailer UF, Frank GK, Henry SE, Price JC, Meltzer CC, Weissfeld L et al. Altered brain serotonin 5-HT1A receptor binding after recovery from anorexia nervosa measured by positron emission tomography and [carbonyl11C]WAY-100635. Arch Gen Psychiatry 2005; 62(9): 1032–1041.

25. Halberstadt AL, Koedood L, Powell SB, Geyer MA. Differential contributions of serotonin receptors to the behavioral effects of indoleamine hallucinogens in mice. J Psychopharmacol 2011; 25(11): 1548–1561.

26. Hartogsohn I. Set and setting, psychedelics and the placebo response: An extra-pharmacological perspective on psychopharmacology. J Psychopharmacol 2016; 30(12): 1259–1267.

27. Aday JS, Heifets BD, Pratscher SD, Bradley E, Rosen R, Woolley JD. Great Expectations: recommendations for improving the methodological rigor of psychedelic clinical trials. Psychopharmacology (Berl) 2022; 239(6): 1989–2010.

28. Wulff AB, Nichols CD, Thompson SM. Preclinical perspectives on the mechanisms underlying the therapeutic actions of psilocybin in psychiatric disorders. Neuropharmacology 2023; 231: 109504.

29. Ly C, Greb AC, Cameron LP, Wong JM, Barragan EV, Wilson PC et al. Psychedelics Promote Structural and Functional Neural Plasticity. Cell Rep 2018; 23(11): 3170–3182.

30. Aleksandrova LR, Phillips AG. Neuroplasticity as a convergent mechanism of ketamine and classical psychedelics. Trends Pharmacol Sci 2021; 42(11): 929–942.

31. Dos Santos RG, Osorio FL, Crippa JA, Riba J, Zuardi AW, Hallak JE. Antidepressive, anxiolytic, and antiaddictive effects of ayahuasca, psilocybin and lysergic acid diethylamide (LSD): a systematic review of clinical trials published in the last 25 years. Ther Adv Psychopharmacol 2016; 6(3): 193–213.

32. Hesselgrave N, Troppoli TA, Wulff AB, Cole AB, Thompson SM. Harnessing psilocybin: antidepressant-like behavioral and synaptic actions of psilocybin are independent of 5-HT2R activation in mice. Proc Natl Acad Sci U S A 2021; 118(17).

33. Jefsen O, Hojgaard K, Christiansen SL, Elfving B, Nutt DJ, Wegener G et al. Psilocybin lacks antidepressant-like effect in the Flinders Sensitive Line rat. Acta Neuropsychiatr 2019; 31(4): 213–219.

34. Meccia J, Lopez J, Bagot RC. Probing the antidepressant potential of psilocybin: integrating insight from human research and animal models towards an understanding of neural circuit mechanisms. Psychopharmacology (Berl) 2023; 240(1): 27–40.

35. Planchez B, Surget A, Belzung C. Animal models of major depression: drawbacks and challenges. J Neural Transm (Vienna) 2019; 126(11): 1383–1408.

36. Liao C, Kwan AC. Applying Reinforcement Learning to Rodent Stress Research. Chronic Stress (Thousand Oaks) 2021; 5: 2470547020984732.

37. Leysen JE, Gommeren W, Janssen PAJ. Identification of non-serotonergic [3H]ketanserin binding sites on human platelets and their role in serotonin release. European Journal of Pharmacology: Molecular Pharmacology 1991; 206(1): 39–45.

38. Pacheco AT, Olson RJ, Garza G, Moghaddam B. Acute psilocybin enhances cognitive flexibility in rats. Neuropsychopharmacology 2023; 48(7): 1011–1020.

39. Madsen MK, Fisher PM, Burmester D, Dyssegaard A, Stenbaek DS, Kristiansen S et al. Psychedelic effects of psilocybin correlate with serotonin 2A receptor occupancy and plasma psilocin levels. Neuropsychopharmacology 2019; 44(7): 1328–1334.

40. Ling S, Ceban F, Lui LMW, Lee Y, Teopiz KM, Rodrigues NB et al. Molecular Mechanisms of Psilocybin and Implications for the Treatment of Depression. CNS Drugs 2022; 36(1): 17–30.

41. Erkizia-Santamaria I, Alles-Pascual R, Horrillo I, Meana JJ, Ortega JE. Serotonin 5-HT(2A), 5-HT(2c) and 5-HT(1A) receptor involvement in the acute effects of psilocybin in mice. In vitro pharmacological profile and modulation of thermoregulation and head-twich response. Biomed Pharmacother 2022; 154: 113612.

42. Odland AU, Kristensen JL, Andreasen JT. Investigating the role of 5-HT2A and 5-HT2C receptor activation in the effects of psilocybin, DOI, and citalopram on marble burying in mice. Behav Brain Res 2021; 401: 113093.

43. Gonzalez-Maeso J, Weisstaub NV, Zhou M, Chan P, Ivic L, Ang R et al. Hallucinogens recruit specific cortical 5-HT(2A) receptor-mediated signaling pathways to affect behavior. Neuron 2007; 53(3): 439–452.

44. Amodeo DA, Hassan O, Klein L, Halberstadt AL, Powell SB. Acute serotonin 2A receptor activation impairs behavioral flexibility in mice. Behav Brain Res 2020; 395: 112861.

45. Odland AU, Kristensen JL, Andreasen JT. The selective 5-HT2A receptor agonist 25CN-NBOH does not affect reversal learning in mice. Behav Pharmacol 2021; 32(5): 448–452.

46. Shao LX, Liao C, Gregg I, Davoudian PA, Savalia NK, Delagarza K et al. Psilocybin induces rapid and persistent growth of dendritic spines in frontal cortex in vivo. Neuron 2021; 109(16): 2535–2544 e2534.

47. Golden CT, Chadderton P. Psilocybin reduces low frequency oscillatory power and neuronal phase-locking in the anterior cingulate cortex of awake rodents. Sci Rep 2022; 12(1): 12702.

48. Depoortere R, Auclair AL, Bardin L, Colpaert FC, Vacher B, Newman-Tancredi A. F15599, a preferential post-synaptic 5-HT1A receptor agonist: activity in models of cognition in comparison with reference 5-HT1A receptor agonists. Eur Neuropsychopharmacol 2010; 20(9): 641–654.

49. Alvarez BD, Morales CA, Amodeo DA. Impact of specific serotonin receptor modulation on behavioral flexibility. Pharmacol Biochem Behav 2021; 209: 173243.

50. Grandjean J, Buehlmann D, Buerge M, Sigrist H, Seifritz E, Vollenweider FX et al. Psilocybin exerts distinct effects on resting state networks associated with serotonin and dopamine in mice. Neuroimage 2021; 225: 117456.

51. Sakashita Y, Abe K, Katagiri N, Kambe T, Saitoh T, Utsunomiya I et al. Effect of psilocin on extracellular dopamine and serotonin levels in the mesoaccumbens and mesocortical pathway in awake rats. Biol Pharm Bull 2015; 38(1): 134–138.

52. den Ouden HE, Daw ND, Fernandez G, Elshout JA, Rijpkema M, Hoogman M et al. Dissociable effects of dopamine and serotonin on reversal learning. Neuron 2013; 80(4): 1090–1100.

53. Foldi CJ, Liknaitzky P, Williams M, Oldfield BJ. Rethinking Therapeutic Strategies for Anorexia Nervosa: Insights From Psychedelic Medicine and Animal Models. Front Neurosci 2020; 14: 43.

54. Attia E, Steinglass JE. Psilocybin for anorexia nervosa: If it helps, let’s learn how. Med 2023; 4(9): 581–582.

55. Maddox WT, Markman AB. The Motivation-Cognition Interface in Learning and Decision Making. Curr Dir Psychol Sci 2010; 19(2): 106–110.

56. Fromer R, Lin H, Dean Wolf CK, Inzlicht M, Shenhav A. Expectations of reward and efficacy guide cognitive control allocation. Nat Commun 2021; 12(1): 1030.

57. Gutierrez E. A rat in the labyrinth of anorexia nervosa: contributions of the activity-based anorexia rodent model to the understanding of anorexia nervosa. Int J Eat Disord 2013; 46(4): 289–301.

58. Milton LK, Oldfield BJ, Foldi CJ. Evaluating anhedonia in the activity-based anorexia (ABA) rat model. Physiol Behav 2018; 194: 324–332.

59. Huang KX, Milton LK, Dempsey H, Power SJ, Conn KA, Andrews ZB et al. Rapid, automated, and experimenter-free touchscreen testing reveals reciprocal interactions between cognitive flexibility and activity-based anorexia in female rats. Elife 2023; 12.

60. Allen PJ, Jimerson DC, Kanarek RB, Kocsis B. Impaired reversal learning in an animal model of anorexia nervosa. Physiol Behav 2017; 179: 313–318.

61. Milton LK, Mirabella PN, Greaves E, Spanswick DC, van den Buuse M, Oldfield BJ et al. Suppression of Corticostriatal Circuit Activity Improves Cognitive Flexibility and Prevents Body Weight Loss in Activity-Based Anorexia in Rats. Biol Psychiatry 2021; 90(12): 819–828.

62. Doss MK, Madden MB, Gaddis A, Nebel MB, Griffiths RR, Mathur BN et al. Models of psychedelic drug action: modulation of cortical-subcortical circuits. Brain 2022; 145(2): 441–456.

63. Anantharaman-Barr HG, Decombaz J. The effect of wheel running and the estrous cycle on energy expenditure in female rats. Physiol Behav 1989; 46(2): 259–263.

64. Verharen JPH, Kentrop J, Vanderschuren L, Adan RAH. Reinforcement learning across the rat estrous cycle. Psychoneuroendocrinology 2019; 100: 27–31.

65. Cora MC, Kooistra L, Travlos G. Vaginal Cytology of the Laboratory Rat and Mouse:Review and Criteria for the Staging of the Estrous Cycle Using Stained Vaginal Smears. Toxicologic Pathology 2015; 43(6): 776–793.

66. Saber I, Milewski A, Reitz AB, Rawls SM, Walker EA. Effects of dopaminergic and serotonergic compounds in rats trained to discriminate a high and a low training dose of the synthetic cathinone mephedrone. Psychopharmacology (Berl) 2019; 236(3): 1015–1029.

67. Van de Kar LD, Javed A, Zhang Y, Serres F, Raap DK, Gray TS. 5-HT2A receptors stimulate ACTH, corticosterone, oxytocin, renin, and prolactin release and activate hypothalamic CRF and oxytocin-expressing cells. J Neurosci 2001; 21(10): 3572–3579.

68. Ichikawa J, Dai J, Meltzer HY. 5-HT(1A) and 5-HT(2A) receptors minimally contribute to clozapine-induced acetylcholine release in rat medial prefrontal cortex. Brain Res 2002; 939(1-2): 34–42.

69. Matikainen-Ankney BA, Earnest T, Ali M, Casey E, Wang JG, Sutton AK et al. An open-source device for measuring food intake and operant behavior in rodent home-cages. Elife 2021; 10.

70. Schneider CA, Rasband WS, Eliceiri KW. NIH Image to ImageJ: 25 years of image analysis. Nat Methods 2012; 9(7): 671–675.

71. Stirling DR, Swain-Bowden MJ, Lucas AM, Carpenter AE, Cimini BA, Goodman A. CellProfiler 4: improvements in speed, utility and usability. BMC Bioinformatics 2021; 22(1): 433.

72. Alaverdashvili M, Hackett MJ, Pickering IJ, Paterson PG. Laminar-specific distribution of zinc: evidence for presence of layer IV in forelimb motor cortex in the rat. Neuroimage 2014; 103: 502–510.

73. Milton LK, Patton T, O’Keeffe M, Oldfield BJ, Foldi CJ. In pursuit of biomarkers for predicting susceptibility to activity-based anorexia in adolescent female rats. Int J Eat Disord 2022; 55(5): 664–677.

74. Peck SK, Shao S, Gruen T, Yang K, Babakanian A, Trim J et al. Psilocybin therapy for females with anorexia nervosa: a phase 1, open-label feasibility study. Nat Med 2023; 29(8): 1947–1953.

75. Fox MA, Stein AR, French HT, Murphy DL. Functional interactions between 5-HT2A and presynaptic 5-HT1A receptor-based responses in mice genetically deficient in the serotonin 5-HT transporter (SERT). Br J Pharmacol 2010; 159(4): 879–887.

76. Smith RL, Barrett RJ, Sanders-Bush E. Neurochemical and behavioral evidence that quipazine-ketanserin discrimination is mediated by serotonin2A receptor. Journal of Pharmacology and Experimental Therapeutics 1995; 275(2): 1050.

77. Keel PK, Dorer DJ, Eddy KT, Franko D, Charatan DL, Herzog DB. Predictors of mortality in eating disorders. Arch Gen Psychiatry 2003; 60(2): 179–183.

78. Button EJ, Chadalavada B, Palmer RL. Mortality and predictors of death in a cohort of patients presenting to an eating disorders service. Int J Eat Disord 2010; 43(5): 387–392.

79. Fadahunsi N, Lund J, Breum AW, Mathiesen CV, Larsen IB, Knudsen GM et al. Acute and long-term effects of psilocybin on energy balance and feeding behavior in mice. Transl Psychiatry 2022; 12(1): 330.

80. Gukasyan N, Davis AK, Barrett FS, Cosimano MP, Sepeda ND, Johnson MW et al. Efficacy and safety of psilocybin-assisted treatment for major depressive disorder: Prospective 12-month follow-up. J Psychopharmacol 2022; 36(2): 151–158.

81. Avena NM, Bocarsly ME. Dysregulation of brain reward systems in eating disorders: neurochemical information from animal models of binge eating, bulimia nervosa, and anorexia nervosa. Neuropharmacology 2012; 63(1): 87–96.

82. Foldi CJ. Taking better advantage of the activity-based anorexia model. Trends in Molecular Medicine 2023.

83. Dalton B, Lewis YD, Bartholdy S, Kekic M, McClelland J, Campbell IC et al. Repetitive transcranial magnetic stimulation treatment in severe, enduring anorexia nervosa: An open longer-term follow-up. Eur Eat Disord Rev 2020; 28(6): 773–781.

84. Ehrlich S, Geisler D, Ritschel F, King JA, Seidel M, Boehm I et al. Elevated cognitive control over reward processing in recovered female patients with anorexia nervosa. J Psychiatry Neurosci 2015; 40(5): 307–315.

85. Santana N, Bortolozzi A, Serrats J, Mengod G, Artigas F. Expression of Serotonin1A and Serotonin2A Receptors in Pyramidal and GABAergic Neurons of the Rat Prefrontal Cortex. Cerebral Cortex 2004; 14(10): 1100–1109.

86. Riad M, Garcia S, Watkins KC, Jodoin N, Doucet E, Langlois X et al. Somatodendritic localization of 5-HT1A and preterminal axonal localization of 5-HT1B serotonin receptors in adult rat brain. J Comp Neurol 2000; 417(2): 181–194.

87. Miquel MC, Doucet E, Riad M, Adrien J, Verge D, Hamon M. Effect of the selective lesion of serotoninergic neurons on the regional distribution of 5-HT1A receptor mRNA in the rat brain. Brain Res Mol Brain Res 1992; 14(4): 357–362.

88. Ferguson SS. Evolving concepts in G protein-coupled receptor endocytosis: the role in receptor desensitization and signaling. Pharmacol Rev 2001; 53(1): 1–24.

89. Hanyaloglu AC, von Zastrow M. Regulation of GPCRs by endocytic membrane trafficking and its potential implications. Annu Rev Pharmacol Toxicol 2008; 48: 537–568.

90. Baldi E, Bucherelli C. The inverted "u-shaped" dose-effect relationships in learning and memory: modulation of arousal and consolidation. Nonlinearity Biol Toxicol Med 2005; 3(1): 9–21.

91. Hogeveen J, Mullins TS, Romero JD, Eversole E, Rogge-Obando K, Mayer AR et al. The neurocomputational bases of explore-exploit decision-making. Neuron 2022; 110(11): 1869–1879 e1865.

92. King MV, Marsden CA, Fone KCF. A role for the 5-HT1A, 5-HT4 and 5-HT6 receptors in learning and memory. Trends Pharmacol Sci 2008; 29(9): 482–492.

93. Tyls F, Palenicek T, Kaderabek L, Lipski M, Kubesova A, Horacek J. Sex differences and serotonergic mechanisms in the behavioural effects of psilocin. Behav Pharmacol 2016; 27(4): 309–320.

94. Effinger DP, Quadir SG, Ramage MC, Cone MG, Herman MA. Sex-specific effects of psychedelic drug exposure on central amygdala reactivity and behavioral responding. Transl Psychiatry 2023; 13(1): 119.

95. Kehagia AA, Murray GK, Robbins TW. Learning and cognitive flexibility: frontostriatal function and monoaminergic modulation. Curr Opin Neurobiol 2010; 20(2): 199–204.

96. Vollenweider FX, Vontobel P, Hell D, Leenders KL. 5-HT modulation of dopamine release in basal ganglia in psilocybin-induced psychosis in man--a PET study with [11C]raclopride. Neuropsychopharmacology 1999; 20(5): 424–433.

97. Sayali C, Barrett FS. The costs and benefits of psychedelics on cognition and mood. Neuron 2023; 111(5): 614–630.

98. Zhou FM, Liang Y, Salas R, Zhang L, De Biasi M, Dani JA. Corelease of dopamine and serotonin from striatal dopamine terminals. Neuron 2005; 46(1): 65–74.

99. Datta MS, Chen Y, Chauhan S, Zhang J, De La Cruz ED, Gong C et al. Whole-brain mapping reveals the divergent impact of ketamine on the dopamine system. Cell Rep 2023: 113491.

100. Rangel-Barajas C, Malik M, Vangveravong S, Mach RH, Luedtke RR. Pharmacological modulation of abnormal involuntary DOI-induced head twitch response in male DBA/2J mice: I. Effects of D2/D3 and D2 dopamine receptor selective compounds. Neuropharmacology 2014; 83: 18–27.

