## Supplementary for "Psilocybin prevents activity-based anorexia in female rats by enhancing cognitive flexibility: contributions from 5-HT1A and 5-HT2A receptor mechanisms"

### Text summary of Supplementary Materials

Contained in this document is all supplementary materials to accompany the manuscript entitled “Psilocybin prevents weight loss in activity-based anorexia by enhancing cognitive flexibility in female rats: contributions from both 5-HT1A and 5-HT2A receptor subtype mechanisms” by Conn K, Milton LK *et al*.

It includes additional methodological information, animal allocation and exclusions, five supplementary figures and full details of all statistical analyses.

### Supplementary Methods

#### Delayed psilocybin administration during activity-based anorexia (ABA)

Two separate cohorts of female Sprague-Dawley rats (7 weeks old) were habituated to running wheels for 7 days and underwent exposure to the ABA paradigm as described in the main methods. To explore the potential that psilocybin could “rescue” weight loss that had already been initiated, rats were administered psilocybin (1.5mg/kg) or saline after 2 days of ABA exposure ($n$=6 saline; $n$=6 psilocybin) and continued throughout the protocol (**Supplementary Figure 7A**). Because the proportion of weight loss exhibited by these rats at the time of administration differed in this cohort, a second group of rats were psilocybin (1.5mg/kg; $n$=6) or saline ($n$=5) at a timepoint tailored to their individual weight loss trajectories (i.e. when they reached 85% of baseline body weight) and subsequently continued throughout the protocol (**Supplementary Figure 7B**). Neither strategy of delayed administration altered weight loss, food intake or running activity (See **Supplementary Figure 7C-G**), and in fact, a larger proportion of control (saline treated) animals were susceptible to ABA in this experiment than is normally observed, suggesting that the stress associated with injections during this critical period of ABA development accelerated weight loss in both treatment groups.

#### RNAscope in situ hybridisation

To visualise 5-HT receptor mRNA, the RNAScope Multiplex Fluorescent Reagent Kit v2 (Advanced Cell Diagnostics, USA) was used according to a modified protocol specified by the manufacturer. To prepare the sections, the slides were baked in a water bath at 60°C for 30 minutes, followed by immersion in 4% paraformaldehyde for 15 minutes at 40°C and dehydration in 50%, 70% and 100% ethanol in distilled water for 5 minutes at room temperature. Sections were then treated with hydrogen peroxide for 10 minutes at room temperature, followed by 5-minute incubation in 1X Target Retrieval Reagent in a steamer with temperature above 90°C and immersion in 100% ethanol. A hydrophobic barrier was drawn around each section using an ImmEdgeTM hydrophobic barrier pen (Vector Laboratories, USA). Sections were treated with protease III for 30 minutes at 40°C, followed by 2-hour incubation with a mixture of probes for 5-HT1AR and 5-HT2AR at 40°C. Signal amplification procedures were performed before mRNA detection with Opal fluorophore dyes diluted in TSA buffer (Opal 520 Reagent Pack, 1:500; Opal 620 Reagent Pack, 1:750. Akoya Biosicences, USA). Finally, sections were incubated with DAPI counterstain for 30 seconds, followed by coverslipping with Vectashield HardSet antifade mounting medium (Vector Laboratories, USA). Slides were stored in the dark at 4°C until imaging.

**Cell profiler analysis pipeline**

Images were pre-processed for analysis using a macro in ImageJ (version 2.3.0, 64 bit), designed to batch process micrographs into a maximum intensity projection for Z-stack compression, converting the images from 16 to 8 bits, and splitting into separate channel files. These split-channel images were then processed with a custom-built pipeline in CellProfiler (version 4.2.1). The pipeline was designed to recognize the primary objects; the nuclei, and the puncta (between 1 to 10 pixels in diameter) representing *Hrt1a* and *Hrt2a* transcripts. The pipeline also created artificial cell membranes around the nuclei (with a radius of 20 pixels), and by doing so, related the transcripts to individual cells. These transcript-expressing cells were only counted if the threshold of a minimum of three transcripts were present within the artificial cell membrane. The number of transcripts were the main factor evaluated by the pipeline, since each punctum represented a singular transcript. It is important to note that neither the size or intensity of the puncta describes the number of transcripts, but instead the number of probes bound to the target mRNA and thus was not statistically analysed.

**Image analysis**

Images were pre-processed for analysis using a custom macro in ImageJ. The macro automatically opened the lif files, created a maximum intensity projection, separate the 3 channels and saved the individual tif files in a new folder. This folder was then fed into a CellProfiler custom pipeline to segment the nuclei using the *IdentifyPrimaryObject* module (typical object diameter 25-200 pixels; thresholding method: Minimum Cross Entropy, smoothing scale: 0). The resulting objects were expanded 20 pixels using the *IdentifySecondaryObjects* module (method: Distance-N) to define the perinuclear cytoplasm where transcripts are expected to be located; these objects were named “Cells” (see **Supplementary Methods Figure A**, nuclei outlined in green, the “cytoplasm” in magenta). The images corresponding to the RNA probes were first processed through the *EnhanceOrSuppressFeatures* module (selected operation: Enhance, feature type: Speckles, feature size: 20). This step enhances the contrast between adjacent spots, enabling a more accurate segregation of adjacent or overlapping puncta. The *IdentifyPrimaryObject* module was applied to the enhanced image (typical object diameter: 1-10 pixels threshold strategy: Adaptive; thresholding method: Otsu). Using the *RelateObjects* module (parent objects: Cells; child objects: Green or Red), we created new object sets corresponding to the cells that contained green or red puncta (see **Supplementary Methods Figure B**, each Cell or Red (puncta) object is initially a unique colour **B’**,**B”**, the relationship between the puncta and the cells is illustrated by the parent Cell object and all its children puncta being the same colour **B”’**). To identify cells expressing both transcripts, a similar procedure was applied by establishing a relationship between RedCell objects (cells containing red puncta, parent) and the Green puncta objects (child) (see **Supplementary Methods Figure C-C”’**). All counts were exported as spreadsheets using the *ExportToSpreadsheet* module.

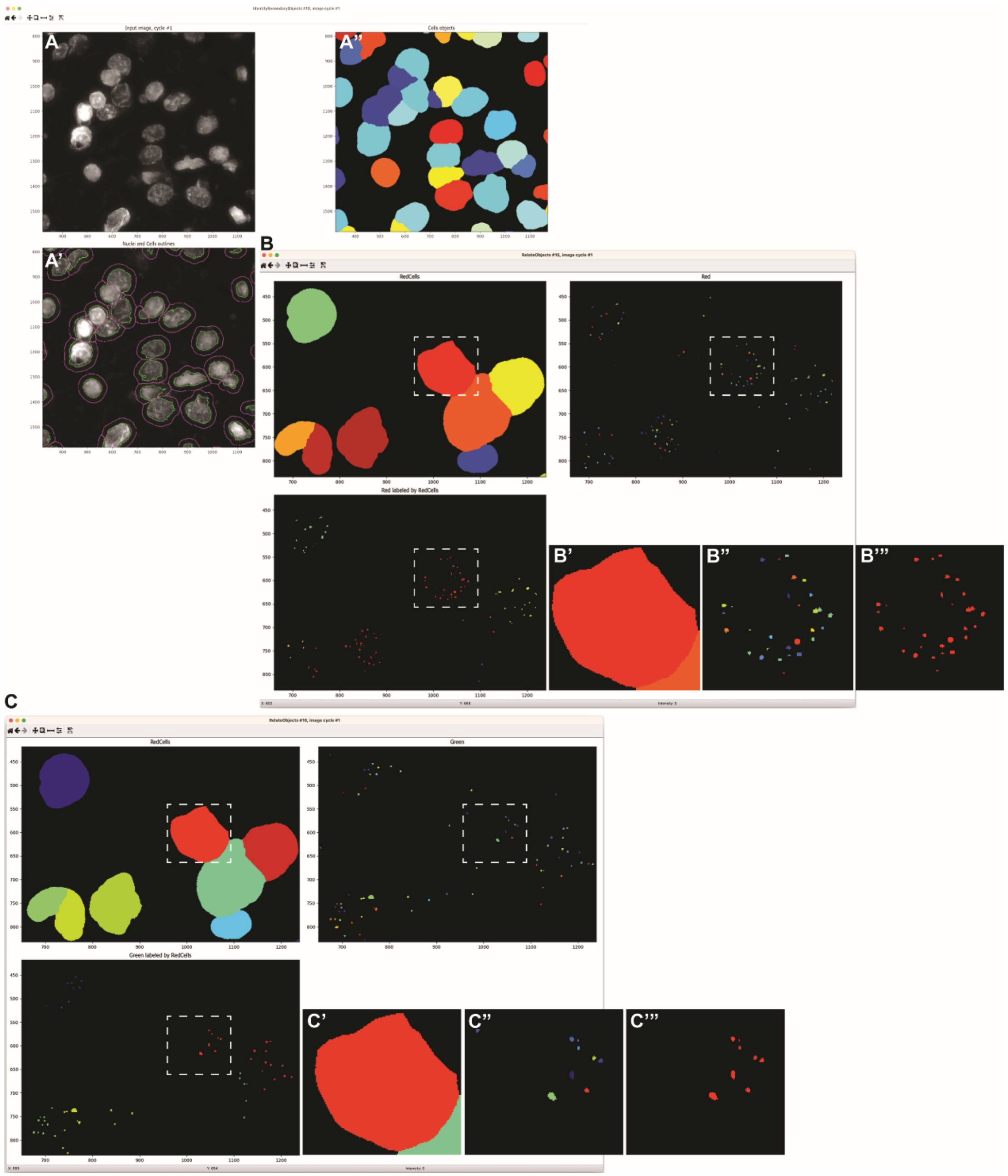

**Supplementary Methods Figure: Segmentation of the nuclei and *Htr1a* (green) and *Htr2a* (red) puncta using a custom CellProfiler pipeline.**

Dapi-labelled nuclei (**A**) were segmented using the minimum cross-entropy thresholding method (**A'**, green outline) and the object was expanded by 20 pixel to account for the perinuclear cytoplasm where transcripts accumulate (**A'**, magenta outline). The resulting Cells objects (parent **A"**) were then related to the puncta objects (children) corresponding to the probes segmented with the Otsu thresholding method, to identify the counts of red or green puncta per cell (**B-B"'**). The cells expressing both transcripts were identified by filtering those positive for one transcript (**C**, RedCells corresponding to the subset of cells expressing *Htr2a*) and establishing a new relationship between RedCells (parent) and Green puncta (children) (**C-C"'**). The same operation was repeated for GreenCells corresponding to the subset of cells expressing *Htr1a)* and also cells negative for one probe.

### Supplementary Table 1. Animal numbers and experimental allocation

| Experiment | Start n | Exclusions | Reasons | Final n | Treatment admin | Figures |
|---|---|---|---|---|---|---|
| Single Dose ABA | 40 | 5 | Didn't adapt to wheel, food hoarding, no RWA data | 35; 16 SAL, 19 PSI | Last day of Baseline;<br>24h before ABA onset | 1, S6 |
| Delayed Dose ABA | 24 | 1 | Food hoarding | 23; 11 SAL 12 PSI | Either after 2 days of ABA or at 85% of baseline body weight | S7 |
| FED between session reversal **SAL+** | 32 | 1 | Didn't learn FR5 | 31; 15 SAL, 16 PSI | Immediately after final FR5 training session;<br>pre-treatment (SAL, KTN, WAY, MDL) given 30 minutes prior to drug (SAL, PSI);<br>18h before Reversal day 1 | 2, 3, S1 |
| FED between session reversal **MDL+** | 28 | 10 | | 18; 9 SAL, 9 PSI | | 3, 4 |
| FED between session reversal **WAY+** | 28 | 2 | | 26; 13 SAL, 13 PSI | | 3, 4, S3 |
| FED between session reversal **KTN+** | 28 | 5 | | 23; 11 SAL, 12 PSI | | S4 |
| Between session reversal **TOTAL** | 116 | 18 | | 98; 48 SAL, 50 PSI | | |
| FED **PR and R-PR** | 28 | 3 | Didn't learn FR5 | 25; 12 SAL, 13 PSI | Immediately after final FR5 training session;<br>18h before PR session | 2, S2 |
| FED Fixed Ratio + **Extinction** | 30 | 8 | | 22; 11 SAL, 11 PSI | Immediately after final FR5 training session;<br>18h before extinction session | 2 |
| FED **Variable Ratio** + **Extinction** | 28 | 5 | Didn't learn FR1 | 23; 11 SAL, 12 PSI | Immediately after final FR1 training session;<br>18h before first VR1-5 session | 2 |
| RNAscope **Time course** | 20 | 1 | Suboptimal detection | 19; 4 SAL, 5 PSI6h, 5 PSI12h, 5 PSI24h | 6, 12, or 24h before brain collection;<br>SAL rats spread across time points;<br>drug admin at same time for all rats, staggered brain collection | 5, S5 |
| RNAscope **anatomy** (SAL and PSI12h from above) | 8 | 0 | | 8, 3 SAL, 5 PSI12h | | 5 |
| RNAscope **Non-ABA** (SAL and PSI6h from above) | 10 | 1 | Suboptimal detection | 9; 4 SAL, 5 PSI6h | | 5, S5 |
| RNAscope **ABA** | 13 | 1 | Suboptimal detection | 12; 5 SAL, 7 PSI | Day that body weight % dropped below 85%;<br>6h before brain collection | 5, S5 |

### Supplementary Figures

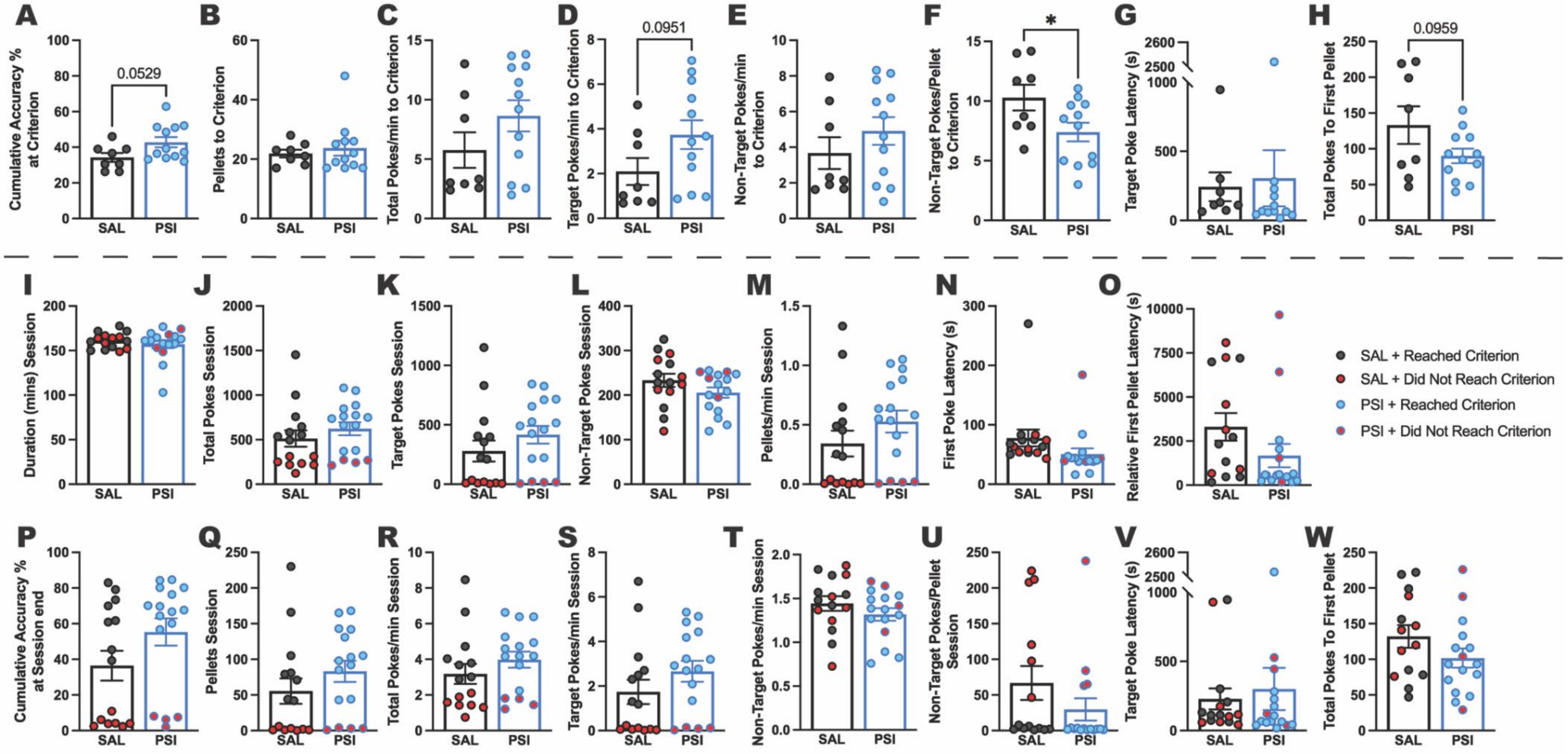

**Supplementary Figure 1. Effects of psilocybin on full session reversal learning measures (including rats that did not meet criterion).** Cumulative accuracy at the point rats reached criterion (80th target poke in a 100-poke moving window) tended to be greater for psilocybin treated rats (**A**, *p*=.0529), although there was no difference in rewards earned at this point (**B**, *p*=.5916). There were no differences in total (**C**, *p*=.1695), target (**D**, *p*=.0951) or non-target (**E**, *p*=.3132) pokes per minute to criterion. Psilocybin treated rats did, however, have a lower rate of non-target pokes per pellet at criterion (**F**, *p*=.0394), while there was no difference in target poke latency (**G**, *p*=.8160), and only a slight trend toward fewer total pokes to earn their first reward (**H**, *p*=.0959). Comparing performance across the entire first reversal session (max 180min; durations in **I** represent time from first to last poke) between the full treatment groups, i.e. all animals regardless of whether they reached criterion (blue/black filled circles) or not (red filled circles), revealed no significant differences between treatment groups on any outcome measure (**I-W**, all *p*s>.1059). Bar graphs show mean ± SEM with individual data points. **p*<.05. SAL saline; PSI psilocybin. For full statistical analysis details see **Supplementary Figure 1 Statistics Table**.

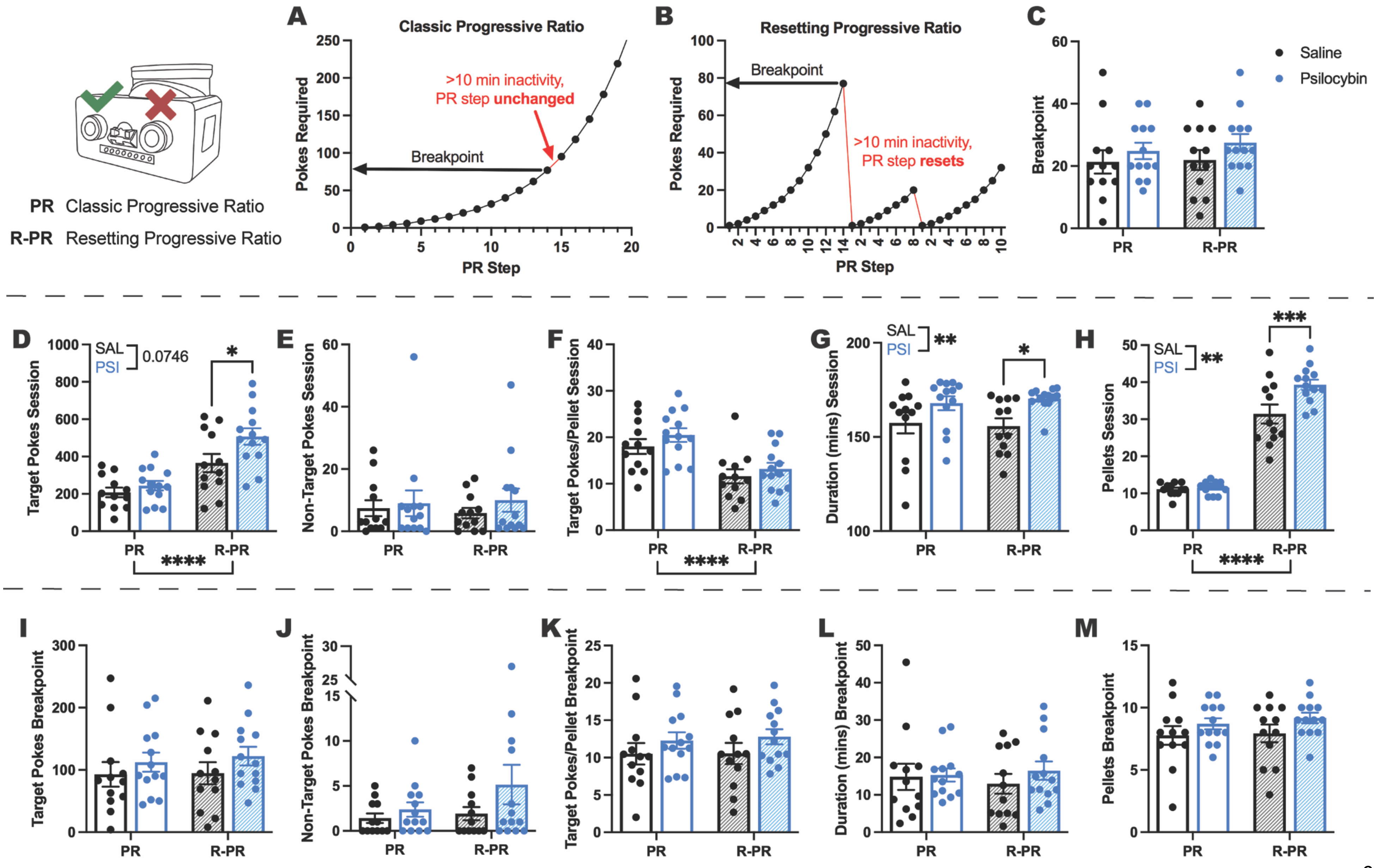
PR Classic Progressive Ratio
R-PR Resetting Progressive Ratio
A
Classic Progressive Ratio
Pokes Required
PR Step
Breakpoint
>10 min inactivity,
PR step unchanged
B
Resetting Progressive Ratio
Pokes Required
PR Step
Breakpoint
>10 min inactivity,
PR step resets
C
Breakpoint
Saline
Psilocybin
PR
R-PR
D
Target Pokes Session
SAL
PSI
0.0746
E
Non-Target Pokes Session
F
Target Pokes/Pellet Session
G
Duration (mins) Session
H
Pellets Session
I
Target Pokes Breakpoint
J
Non-Target Pokes Breakpoint
K
Target Pokes/Pellet Breakpoint
L
Duration (mins) Breakpoint
M
Pellets Breakpoint

**Supplementary Figure 2. Effects of psilocybin on task engagement during a progressive ratio “resetting” task.** On separate days, with a FR5 reinstatement training session between (data not shown) rats underwent a classic progressive ratio (PR) session (**A**, using the formula (5 * e(0.2*$n$)- 5), where $n$ is the trial number) and a resetting progressive ratio (R-PR) session (**B**) where any 10-minute period of inactivity caused the PR step to reset. The classical measure of motivation, breakpoint (**C**), i.e. number of pokes made to earn the last pellet obtained before a 10-minute period of inactivity, along with more specific outcome measures up to this time point of the session (**I-M**) did not differ between schedules or treatment groups, nor was there any interaction between the two (all $p$s>.1089), indicating that up to breakpoint (10 minutes of inactivity) both tasks were conceptually ‘analogous’ producing similar performance that was not effected by psilocybin treatment. Conversely, analysis of entire sessions (max 180min), total target pokes (**D**, Schedule $p<.0001$, Treatment $p=.0746$, Interaction $p=.0346$), session duration (time from first to last poke; **G**, Schedule $p=.9299$, Treatment $p=.0053$, Interaction $p=.5919$), and pellets earned (**H**, Schedule $p<.0001$, Treatment $p=.0058$, Interaction $p=.0260$) revealed significant differences, whereas non-target pokes (**E**, all $p$s>.3983) did not, and target pokes/pellet revealed only an expected effect of schedule (**F**, Schedule $p<.0001$, Treatment $p=.3251$, Interaction $p=.5073$) inherent to the resetting nature of the R-PR schedule. Importantly, all treatment group differences were restricted to the R-PR session and occurred in the same direction, with psilocybin treated rats executing more target pokes (**D**, R-PR SAL<PSI $p=.0209$), over a longer period of task engagement (**G**, R-PR SAL<PSI $p=.0235$), culminating in a greater number of pellets earned (**H**, R-PR SAL<PSI $p=.0009$) than saline treated rats. Bar graphs show mean ± SEM with individual data points. *$p<.05$, **$p<.01$, ***$p<.001$, ****$p<.0001$. PR classic progressive ratio; R-PR resetting progressive ratio; SAL saline; PSI psilocybin. For full statistical analysis details see **Supplementary Figure 2 Statistics Table**.

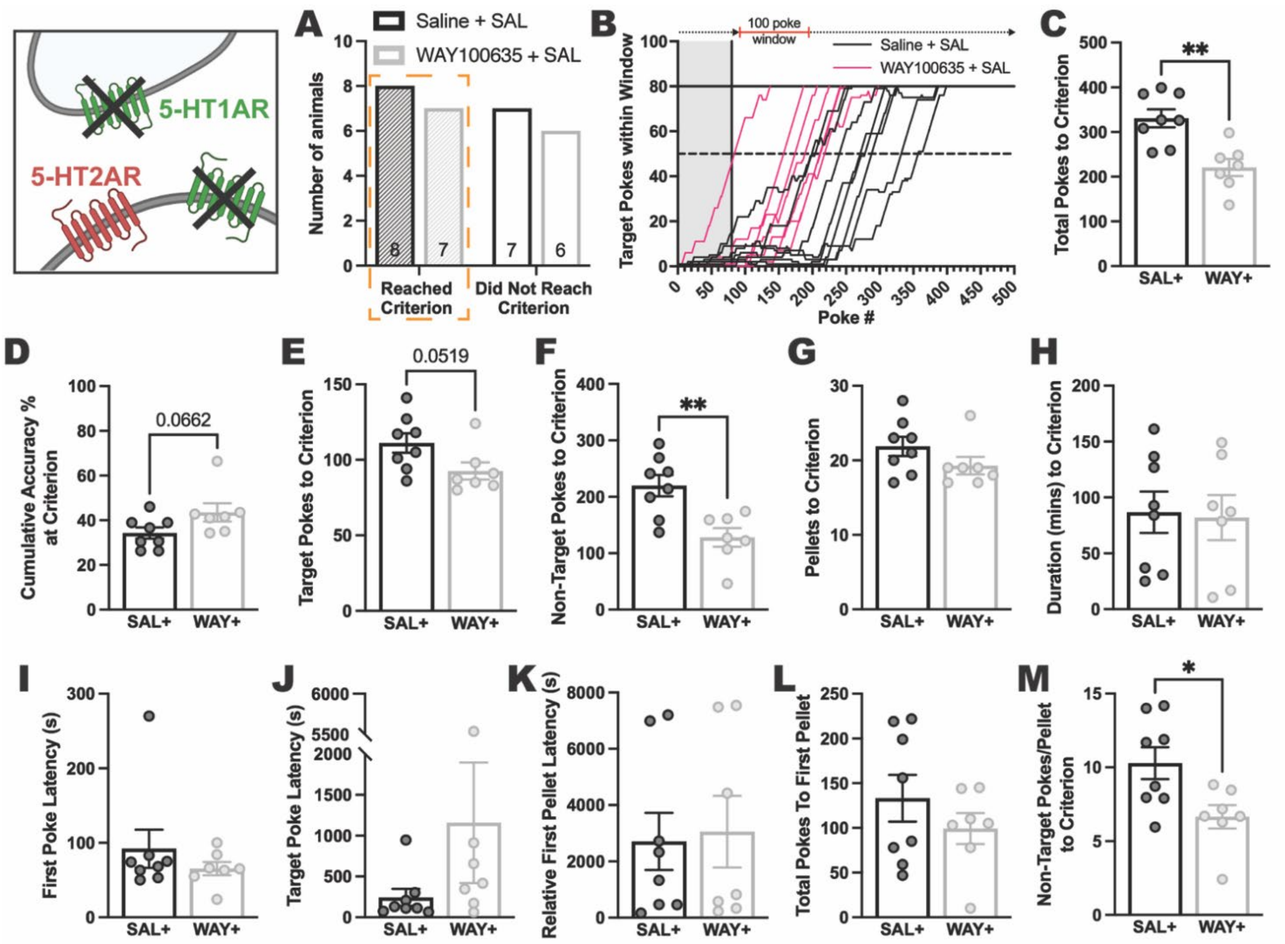

**Supplementary Figure 3. Effects of 5-HT1AR antagonism on reversal learning performance to criterion in control rats.** Compared to control rats (Saline+SAL), antagonism of 5-HT1AR (WAY+SAL, henceforth WAY+) produced a near identical split of rats reaching the reversal day 1 performance criterion (**A**) of 80 target pokes within a 100-poke moving window (**B**) in significantly fewer total pokes (**C**, *p*=.0017). There was a trend toward increased accuracy (**D**, *p*=.0662) and greater target pokes (**E**, *p*=.0519), accompanied by significantly fewer non-target pokes (**F**, *p*=.0033), although the groups obtained a similar number of pellets (**G**, *p*=.1650) and reached criterion in a similar time frame (time from first poke to poke that achieved criterion; **H**, *p*=.8615). There was no effect of 5-HT1AR antagonism on first poke (time from device access to first poke; **I**, *p*=.3717), target poke (**J**, *p*=.2114), nor relative first pellet (time from first poke to earning first pellet; **K**, *p*=.8310) latency. While both groups required a similar number of total pokes to acquire their first pellet (**L**, *p*=.3159), WAY+ had significantly lower non-target pokes/pellet at criterion (**M**, *p*=.0206). Bar graphs show mean ± SEM with individual data points. **p*<.05, ***p*<.01. SAL saline; SAL+ Saline+SAL; WAY+ WAY100635+SAL. For full statistical analysis details see **Supplementary Figure 3 Statistics Table**.

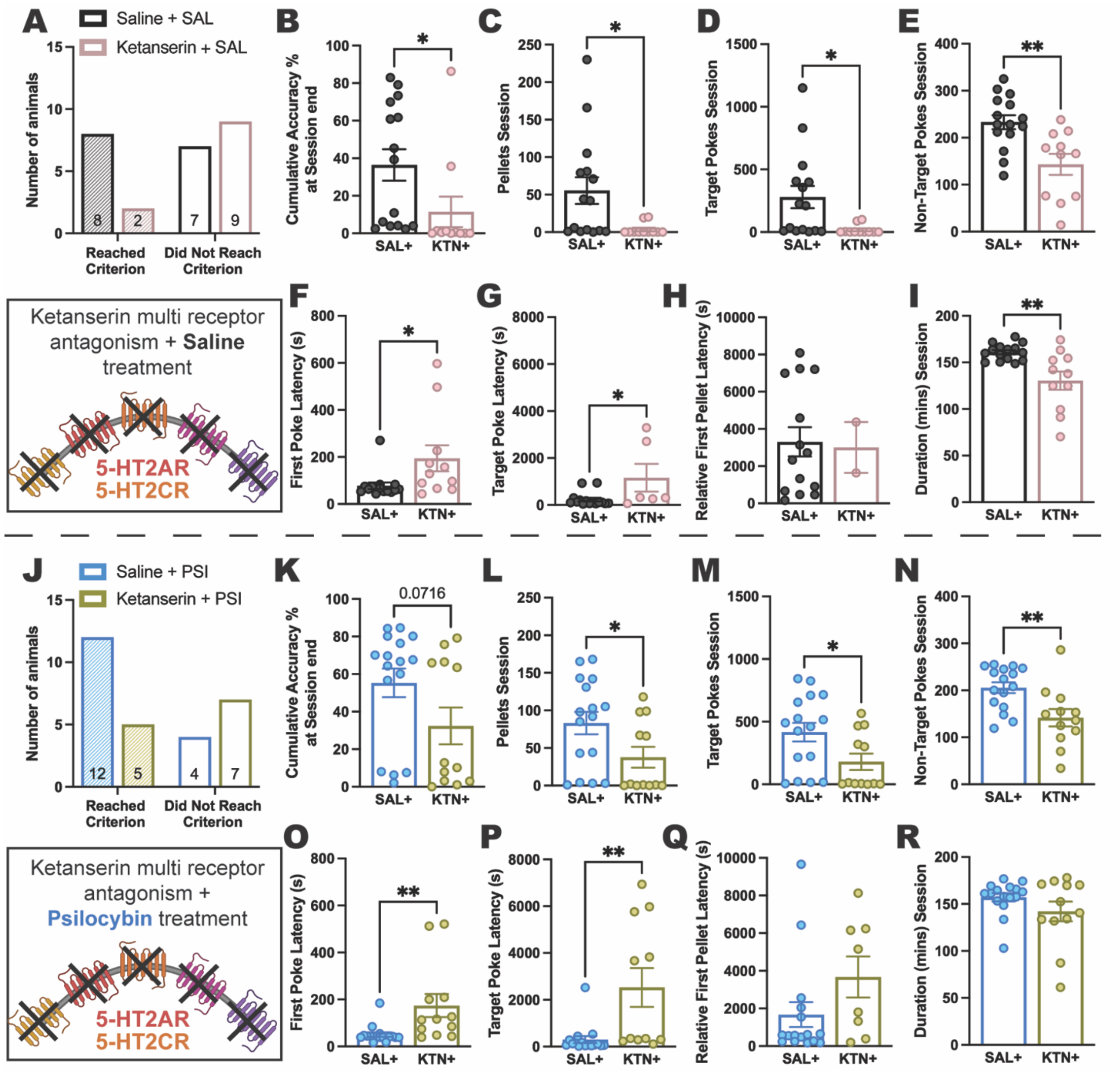
A
Saline + SAL
Ketanserin + SAL
Number of animals
Reached Criterion
Did Not Reach Criterion
B
Cumulative Accuracy % at Session end
C
Pellets Session
D
Target Pokes Session
E
Non-Target Pokes Session
SAL+
KTN+
Ketanserin multi receptor antagonism + **Saline** treatment
5-HT2AR
5-HT2CR
F
First Poke Latency (s)
G
Target Poke Latency (s)
H
Relative First Pellet Latency (s)
I
Duration (mins) Session
J
Saline + PSI
Ketanserin + PSI
K
0.0716
L
M
N
Ketanserin multi receptor antagonism + **Psilocybin** treatment
O
P
Q
R

**Supplementary Figure 4: Effects of ketanserin on reversal learning performance in control and psilocybin treated rats.** Reversal learning in saline treated animals following multi receptor antagonism via pre-treatment with ketanserin (**A**; KTN+) was impaired with only 2/11 (18.2%) rats reaching criterion, compared to 8/15 (53.3%) of control (SAL+) rats. KTN+ produced deficits across the board, resulting in decreased accuracy (**B**, $p$=.0491), fewer pellets earned (**C**, $p$=.0212), target pokes (**D**, $p$=.0200) and non-target pokes (**E**, $p$=.0019), and delaying both first poke (time from device access to first poke; **F**, $p$=.0284) and first target poke (**G**, $p$=.0233) latencies, with only 6/9 (66.7%) KTN+ rats making a target poke, while only 2/9 (22.2%) KTN+ rats earned a pellet (i.e. made at least 5 target pokes; **H**, time from first poke to earning first pellet). KTN+ rats also engaged with the task for a shorter duration (time from first to last poke; **I**, $p$=.0018). A more moderate impairment was seen in psilocybin treated rats following KTN+ pre-treatment, with 5/12 (41.7%) rats reaching criterion compared to 12/16 (75%) Saline+PSI rats (**J**). Although KTN+PSI only showed a trend toward lower accuracy (**K**, $p$=.0716), they earned significantly fewer pellets (**L**, $p$=.0397), made significantly fewer target (**M**, $p$=.0306), and non-target (**N**, $p$=.0048) pokes, as well as exhibiting delayed first poke (**O**, $p$=.0088) and target poke (**P**, $p$=.0041) latencies, although 11/12 (91.7%) rats made a target poke. There was no difference between psilocybin treated groups for relative first pellet latency (**Q**, $p$=.1128, 8/12 [66.7%] KTN+ earned a pallet), nor time spent engaging with the task (**R**, $p$=.1592). Bar graphs show mean ± SEM with individual data points. *$p$<.05, **$p$<.01. SAL saline; PSI psilocybin; SAL+ Saline+SAL or Saline+PSI; KTN+ Ketanserin+SAL or Ketanserin+PSI. For full statistical analysis details see **Supplementary Figure 4 Statistics Table**.

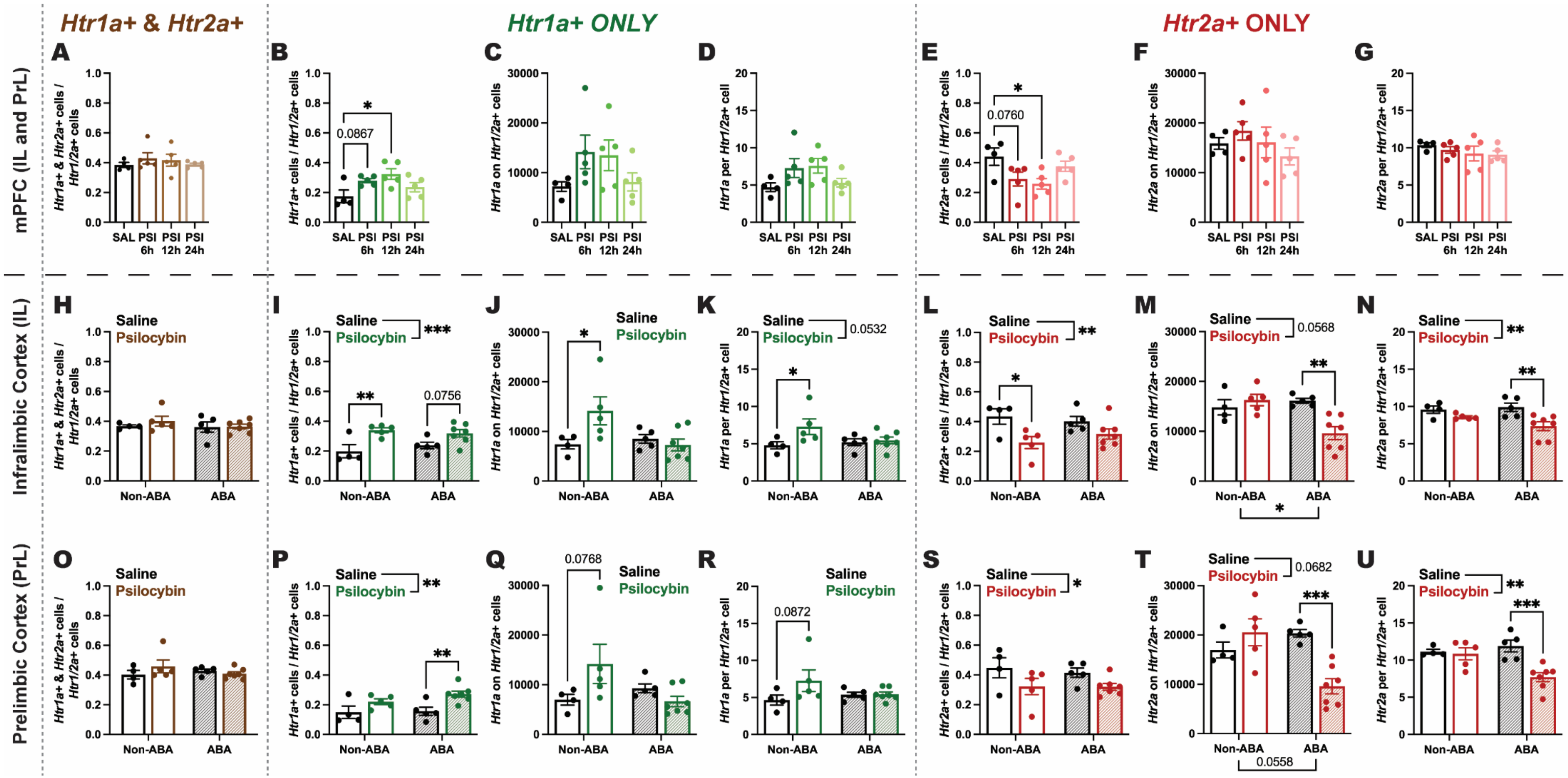

**Supplementary Figure 5. Additional effects of psilocybin on *Htr1a* and *Htr2a* transcript abundance.** The proportion of mPFC *Htr1/2a*+ cells that were double labelled (**brown**) with *Htr1a* and *Htr2a* was not changed by psilocybin treatment (**A**, $p=.6785$). Psilocybin treatment changed the proportion of exclusively *Htr1a* labelled (***Htr1a* ONLY; green**) mPFC *Htr1/2a*+ cells (**B**, $p=.0312$) trending to an increase 6h post administration (SAL<PSI6h $p=.0867$) that was significant 12h post administration (SAL<PSI12h $p=.0137$), although there was no corresponding effect in the absolute number of *Htr1a* transcripts (**C**, $p=.1859$) nor *Htr1a* copies per mPFC *Htr1/2a*+ cell (**D**, $p=.1204$). Psilocybin had the opposite effect on the proportion of exclusively *Htr2a* labelled cells (***Htr2a* ONLY; red**) at the same time points (**E**, $p=.0445$, SAL>PSI6h $p=.0760$, SAL>PSI12h $p=.0295$), with the same absence of effect in absolute copies (**F**, $p=.4230$) and copies per cell (**G**, $p=.5509$). Neither psilocybin treatment nor ABA exposure altered the proportion of *Htr1a* and *Htr2a* double labelled cells in either the IL (**H**, all $ps>.4193$) or PrL (**O**, all $ps>.1597$). ABA exposure had no overall effect on *Htr1a* expression in the IL (**I-K**, ABA exposure $ps>.2032$) or the PrL (**P-R**, ABA exposure $ps>.2329$). Psilocybin significantly increased the proportion of exclusively *Htr1a* labelled IL *Htr1/2a*+ cells (**I**, Treatment $p=.0008$), and produced a near significant increase in *Htr1a* copies per cell (**K**, Treatment *p*=.0532), with no effect on absolute *Htr1a* transcript number (**J**, Treatment *p*=.1202). Although there was only a significant interaction of psilocybin with ABA exposure for this latter measure (**J**, Interaction *p*=.0302), psilocybin significantly increased all outcome measures compared to saline within only the Non-ABA group (**I**, Non-ABA SAL<PSI *p*=.0068; **J**, Non-ABA SAL<PSI *p*=.0333; **K**, Non-ABA SAL<PSI *p*=.0446). ABA exposure had a significant overall effect (and interaction with psilocybin treatment) on the number of *Htr2a* transcripts on IL *Htr1/2a*+ cells (**M**, ABA exposure *p*=.0458, Interaction *p*=.0050), with no effect (nor treatment interaction) for the proportion of exclusively *Htr2a* labelled cells (**L**, ABA exposure *p*=.7620, Interaction *p*=.2769) nor *Htr2a* copies per cell (**N**, ABA exposure *p*=.4118, Interaction *p*=.1663). Psilocybin treatment significantly reduced both the proportion of exclusively *Htr2a* labelled IL *Htr1/2a*+ cells (**L**, Treatment *p*=.0047) and *Htr2a* copies per cell (**N**, Treatment *p*=.0042), with a near significant decrease in the absolute number of *Htr2a* transcripts (**M**, Treatment *p*=.0568). Psilocybin significantly reduced the proportion of exclusively *Htr2a* labelled IL *Htr1/2a*+ cells only in the Non-ABA group (**L**, Non-Aba SAL>PSI *p*=.0197), whereas for both the absolute number of *Htr2a* transcripts (**M**, ABA SAL>PSI *p*=.0018) and *Htr2a* copies per cell (**N**, ABA SAL>PSI *p*=.0043) a significant reduction following psilocybin was found only in the ABA exposed group. Psilocybin significantly increased the overall proportion of exclusively *Htr1a* labelled PrL *Htr1/2a*+ cells (**P**, Treatment *p*=.0029), with no effect on the absolute number of *Htr1a* copies (**Q**, Treatment *p*=.2989) or *Htr1a* copies per cell (**R**, Treatment *p*=.1107), and a significant interaction with ABA exposure only for absolute number of *Htr1a* copies (**Q**, Interaction *p*=.0337). Psilocybin significantly increased the proportion of exclusively *Htr1a* labelled PrL *Htr1/2a*+ cells in the ABA exposed group (**P**, ABA SAL<PSI *p*=.0088), with trends towards increasing both the absolute number of *Htr1a* copies (**Q**, Non-ABA SAL<PSI *p*=.0768) and *Htr1a* copies per cell (**R**, Non-ABA SAL<PSI *p*=.0872) this time in the Non-ABA exposed group. ABA exposure had a near significant effect on the absolute number of *Htr2a* copies on PrL *Htr1/2a*+ cells (**T**, ABA exposure *p*=.0558), but did not change the proportion of exclusively *Htr2a* labelled cells (**S**, ABA exposure *p*=.6905), nor *Htr2a* copies per cell (**U**, ABA exposure *p*=.1091). Psilocybin significantly decreased the proportion of exclusively *Htr2a* labelled PrL *Htr1/2a*+ cells (**S**, Treatment *p*=.0189) and *Htr2a* copies per cell (**U**, Treatment *p*=.0057), with a trend level effect on the absolute number of *Htr2a* transcripts (**T**, Treatment *p*=.0682). There was a significant interaction between psilocybin and ABA exposure for both absolute *Htr2a* transcript number (**T**, Interaction *p*=.0012) and *Htr2a* copies per cell (**U**, Interaction *p*=.0135), resulting in a strong reduction in both measures exclusively within the ABA exposed group (**T**, ABA SAL>PSI *p*=.0007; **U**, ABA SAL>PSI *p*=.0006). Data show mean ± SEM with individual data points. Values are the average of 4 (PrL and IL) or 8 (mPFC) sections per animal. **p*<.05, ***p*<.01, ****p*<.001. SAL saline; PSI psilocybin; PrL prelimbic cortex; IL infralimbic cortex; mPFC medial prefrontal cortex (PrL and IL combined); *Htr1/2a*+ cells expressing *Htr1a* and/or *Htr2a*; ABA activity-based anorexia. For full statistical analysis details see **Supplementary Figure 5 Statistics Table**.

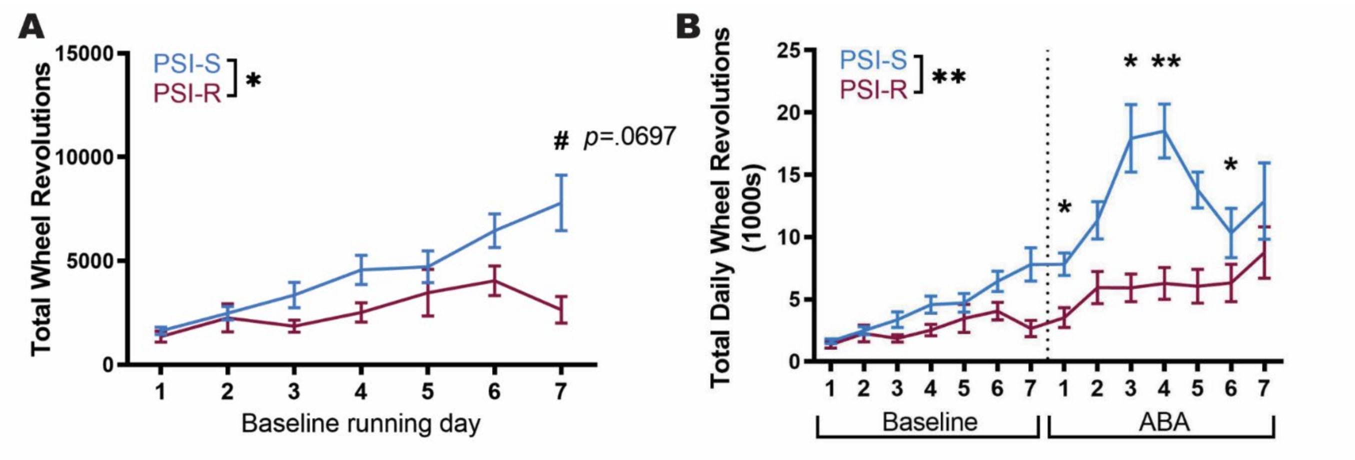

**Supplementary Figure 6. Baseline running prior to administration of psilocybin predicts response outcomes.** Psilocybin treated animals that responded to treatment with positive body weight outcomes (PSI-R) already engaged in less wheel running at baseline (**A**; Day $p<.0001$; ABA Outcome $p=.0221$; Interaction $p=.0071$) than non-responders (PSI-S), although no individual timepoint was significantly different after correcting for multiple comparisons. We also statistically analysed the data presented in **Fig 1K** using a mixed-model ANOVA to demonstrate some specific timepoints during ABA are statistically different between responders and non-responders (**B**), however it should be noted that because of the nature of the ABA model that requires rats (therefore data points) to be removed over successive days of exposure, these statistics could be misleading. Thus, we would point the reader to the reliable analysis in **Fig 1L** that accounts for the variation in number of days each animal remained under ABA conditions. PSI-R, psilocybin resistant; PSI-S, psilocybin susceptible; ABA, activity-based anorexia. For full statistical analysis details see **Supplementary Figure 6 Statistics Table**.

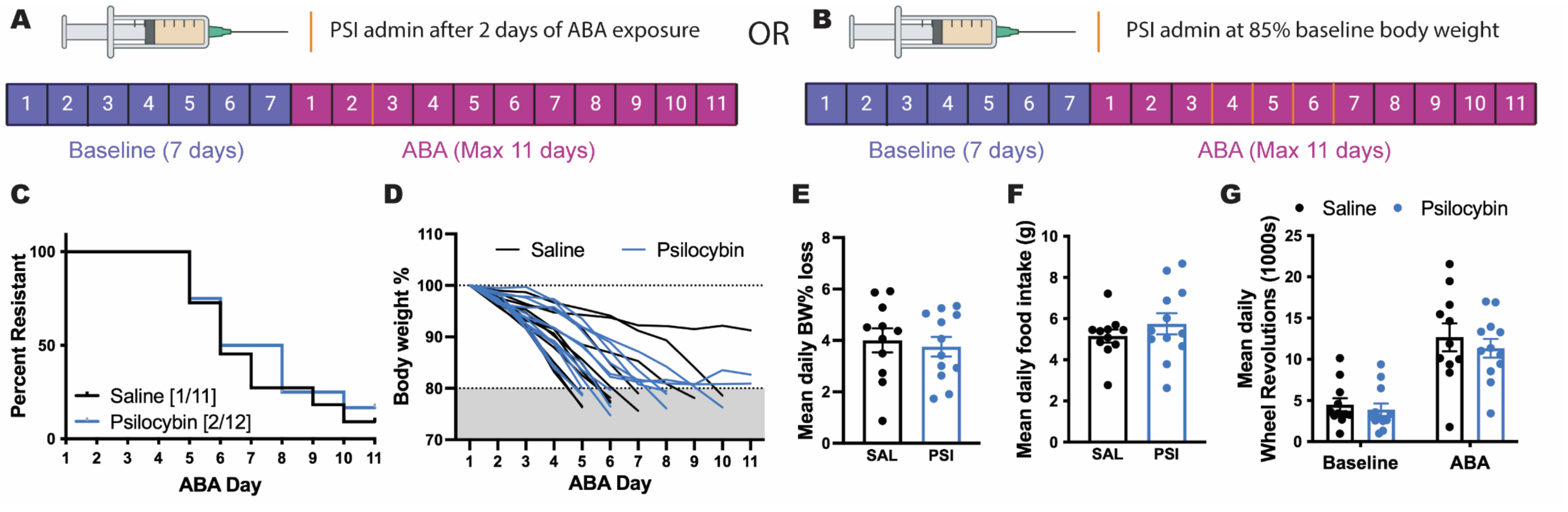

**Supplementary Figure 7. Effects of delayed administration of psilocybin on rescue of ABA phenotype.** Animals were administered psilocybin or saline after weight loss in ABA had initiated, either after a time-based delay of 2 days (**A**) or a weight loss-based delay of 85% baseline body weight (**B**). Orange lines in the timeline figure indicate administration time points. Only 1 saline treated (9%) and 2 psilocybin treated (16%) animal was resistant to weight loss in ABA (**C**) and individual weight loss trajectories were similar between groups (**D**). Delayed psilocybin treatment did not significantly alter daily weight loss on average (**E**; $p$=.6802), daily food intake (**F**; $p$=.3444) and while both groups showed the expected increase in running from baseline (**G P**hase; $p$=.0001) average running wheel activity did not differ between treatment groups (**G** Treatment; $p$=.5261). Bar graphs show mean ± SEM with individual data points for both delayed administration paradigms combined into the same figures. SAL saline; PSI psilocybin; ABA activity-based anorexia. For full statistical analysis details see **Supplementary Figure 7 Statistics Table**.

### Statistics Table - Figure 1

| Figure | Statistical Test | Group n | Main analysis result | Post-hoc multiple comparisons of interest |
|---|---|---|---|---|
| **1C** | Unpaired t-test | SAL n=16;<br>PSI n=19 | $t$(33)=2.508, **$p$=.0172** | |
| **1D** | | | $t$(33)=1.918, $p$=.0638 | |
| **1E** | | | $t$(33)=2.146, **$p$=.0394** | |
| **1G** | RM Two-way ANOVA;<br>Bonferroni's MC | | ABA Phase $F$(1, 33)=126.5, **$p$<.0001**<br>Treatment $F$(1, 33)=1.159, $p$=.2895<br>Interaction $F$(1, 33)=1.033, $p$=.3169 | Baseline: SAL vs PSI $p$>.9999<br>ABA: SAL vs PSI $p$=.3089 |
| **1H** | Unpaired t-test | | $t$(33)=1.098, $p$=.2800 | |
| **1J** | | | $t$(33)=0.9908, $p$=.3290 | |
| **1L** | RM Two-way ANOVA;<br>Bonferroni's MC | PSI-S n=11;<br>PSI-R n=8 | ABA Phase $F$(1, 17)=36.48, **$p$<.0001**<br>ABA Outcome $F$(1, 17)=19.01, **$p$<.0001**<br>Interaction $F$(1, 17)=17.09, **$p$=.0047** | Baseline: PSI-S vs PSI-R $p$=.7415<br>ABA: PSI-S > PSI-R **$p$<.0001** |
| **1M** | Unpaired t-test | | $t$(17)=6.203, **$p$<.0001** | |
| **1O** | | | $t$(17)=2.577, **$p$=.0196** | |
| **1Q** | RM Two-way ANOVA;<br>Bonferroni's MC | SAL-S<br>n=14; PSI-S<br>n=11 | ABA Phase $F$(1, 24)=126.5, **$p$<.0001**<br>Treatment $F$(1, 24)=1.159, $p$=.2895<br>Interaction $F$(1, 24)=1.033, $p$=.3169 | |
| **1R** | Unpaired t-test | | $t$(24)=0.6452, $p$=.5252 | |
| **1T** | | | $t$(24)=0.4783, $p$=.6368 | |

### Statistics Table - Figure 2

| Figure | Statistical Test | Group n | Main analysis result |
|---|---|---|---|
| **2B** | Mixed-effects model<br>**ONLY on Reversal Day 1,**<br>6 x 30min time bins | SAL n=15; PSI n=16 | Time bin $F(2.527, 64.69)=20.04$, **$p<.0001$**<br>Treatment $F(1, 29)=5.128$, **$p=.0312$**<br>Interaction $F(5, 128)=1.530$, $p=.1850$ |
| **2E** | Unpaired t-test | SAL n=8; PSI n=12 | $t(18)=1.514$, $p=.1474$ |
| **2F** | | | $t(18)=1.536$, $p=.1420$ |
| **2G** | | | $t(18)=0.5614$, $p=.5815$ |
| **2H** | | | $t(18)=2.425$, **$p=.0260$** |
| **2I** | | | $t(18)=1.759$, $p=.0956$ |
| **2J** | | | $t(18)=2.307$, **$p=.0332$** |
| **2K** | | | $t(18)=2.291$, **$p=.0343$** |
| **2L** | | SAL n=12; PSI n=13 | $t(23)=0.7795$, $p=.4436$ |
| **2M** | RM Two-way ANOVA | SAL n=9; PSI n=10 | Time bin $F(1.531, 26.02)=42.79$, **$p<.0001$**<br>Treatment $F(1, 17)=0.3212$, $p=.5783$<br>Interaction $F(179, 3043)=0.2623$, $p>.9999$ |
| **2N** | | SAL n=11; PSI n=12 | Schedule $F(2, 42)=18.45$, **$p<.0001$**<br>Treatment $F(1, 21)=1.741$, $p=.2013$<br>Interaction $F(2, 42)=0.7244$, $p=.4906$ |
| **2O** | | | Time bin $F(1.633, 34.28)=22.60$, **$p<.0001$**<br>Treatment $F(1, 21)=0.2681$, $p=.6100$<br>Interaction $F(179, 3759)=0.3509$, $p>.9999$ |

### Statistics Table - Figure 3

| Figure | Statistical Test | Group n | Main analysis result | Post-hoc multiple comparisons of interest |
|---|---|---|---|---|
| **3D** | One-way ANOVA; Dunnett's MC comparing MDL+ and WAY+ to SAL+ | SAL+SAL n=15;<br>MDL+SAL n=9;<br>WAY+SAL n=13 | *F*(2, 34)=4.896, ***p*=.0135** | SAL+ > MDL+ ***p*=.0221**, SAL+ vs WAY+ *p*=.9096 |
| **3E** | | | *F*(2, 34)=3.230, *p*=.0520 | SAL+ > MDL+ ***p*=.0324**, SAL+ vs WAY+ *p*=.7347 |
| **3F** | | | *F*(2, 34)=3.303, ***p*=.0489** | SAL+ > MDL+ ***p*=.0292**, SAL+ vs WAY+ *p*=.6621 |
| **3G** | | | *F*(2, 34)=7.664, ***p*=.0018** | SAL+ > MDL+ ***p*=.0048**, SAL+ > WAY+ ***p*=.0041** |
| **3H** | | | *F*(2, 34)=1.172, *p*=.3218 | SAL+ vs MDL+ *p*=.4572, SAL+ vs WAY+ *p*=.8359 |
| **3I** | | SAL+SAL n=15;<br>MDL+SAL n=6;<br>WAY+SAL n=11 | *F*(2, 29)=4.671, ***p*=.0174** | SAL+ < MDL+ *p*=.0502, SAL+ < WAY+ ***p*=.0244** |
| **3J** | | SAL+SAL n=14;<br>MDL+SAL n=3;<br>WAY+SAL n=9 | *F*(2, 23)=1.797, *p*=.1883 | SAL+ vs MDL+ *p*=.1507, SAL+ vs WAY+ *p*=.9997 |
| **3K** | | SAL+SAL n=15;<br>MDL+SAL n=9;<br>WAY+SAL n=13 | *F*(2, 34)=0.5772, *p*=.5669 | SAL+ vs MDL+ *p*=.6048, SAL+ vs WAY+ *p*=.5604 |
| **3O** | | SAL+PSI n=16;<br>MDL+ PSI n=9;<br>WAY+PSI n=13 | *F*(2, 35)=6.196, ***p*=.0050** | SAL+ vs MDL+ *p*=.2837, SAL+ > WAY+ ***p*=.0024** |
| **3P** | | | *F*(2, 35)=7.089, ***p*=.0026** | SAL+ > MDL+ *p*=.0698, SAL+ > WAY+ ***p*=.0015** |
| **3Q** | | | *F*(2, 35)=7.364, ***p*=.0021** | SAL+ > MDL+ *p*=.0542, SAL+ > WAY+ ***p*=.0013** |
| **3R** | | | *F*(2, 35)=3.995, ***p*=.0274** | SAL+ > MDL+ ***p*=.0210**, SAL+ vs WAY+ *p*=.9497 |
| **3S** | | | *F*(2, 35)=1.586, *p*=.2191 | SAL+ vs MDL+ *p*=.4886, SAL+ vs WAY+ *p*=.1636 |
| **3T** | | SAL+PSI n=16;<br>MDL+ PSI n=7;<br>WAY+PSI n=10 | *F*(2, 30)=3.804, ***p*=.0337** | SAL+ vs MDL+ *p*=.2991, SAL+ < WAY+ ***p*=.0212** |
| **3U** | | SAL+PSI n=16;<br>MDL+PSI n=5;<br>WAY+PSI n=7 | *F*(2, 25)=1.371, *p*=.2724 | SAL+ vs MDL+ *p*=.4930, SAL+ vs WAY+ *p*=.2588 |
| **3V** | | SAL+PSI n=16;<br>MDL+PSI n=9;<br>WAY+PSI n=13 | *F*(2, 35)=1.423, *p*=.2547 | SAL+ vs MDL+ *p*=.4345, SAL+ vs WAY+ *p*=.7309 |

### Statistics Table - Figure 4

| Figure | Statistical Test | Group n | Main analysis result |
|---|---|---|---|
| **4B** | Unpaired t-test | MDL+SAL n=9; MDL+PSI n=9 | *t*(16)=3.034, ***p*=.0079** |
| **4C** | | | *t*(16)=2.255, ***p*=.0385** |
| **4D** | | | *t*(16)=2.191, ***p*=.0436** |
| **4E** | | | *t*(16)=0.7609, *p*=.4578 |
| **4F** | | | *t*(16)=1.034, *p*=.3163 |
| **4G** | | MDL+SAL n=6; MDL+PSI n=7 | *t*(11)=0.5098, *p*=.6202 |
| **4H** | | MDL+SAL n=3; MDL+PSI n=5 | *t*(6)=1.295, *p*=.2428 |
| **4I** | | MDL+SAL n=9; MDL+PSI n=9 | *t*(16)=1.133, *p*=.2741 |
| **4K** | | WAY+SAL n=13; WAY+PSI n=13 | *t*(24)=2.011, *p*=.0556 |
| **4L** | | | *t*(24)=1.806, *p*=.0834 |
| **4M** | | | *t*(24)=1.771, *p*=.0893 |
| **4N** | | | *t*(24)=1.317, *p*=.2001 |
| **4O** | | | *t*(24)=0.7925, *p*=.4358 |
| **4P** | | WAY+SAL n=11; WAY+PSI n=10 | *t*(19)=0.1996, *p*=.8440 |
| **4Q** | | WAY+SAL n=9; WAY+PSI n=7 | *t*(14)=0.3062, *p*=.7639 |
| **4R** | | WAY+SAL n=13; WAY+PSI n=13 | *t*(24)=0.1337, *p*=.8948 |

### Statistics Table - Figure 5

| Figure | Statistical Test | Group n | Main analysis result | Post-hoc multiple comparisons of interest |
|---|---|---|---|---|
| **5B** | One-way ANOVA; Dunnett's MC comparing all PSI groups to SAL | SAL n=4; PSI6h n=5; PSI12h n=5; PSI24h n=5 | $F(3, 15)=0.7801$, $p=.5233$ | |
| **5C** | | | $F(3, 15)=0.2449$, $p=.8637$ | |
| **5D** | | | $F(3, 15)=2.443$, $p=.1043$ | SAL vs PSI6h $p=.2876$, SAL < PSI12h **$p=.0500$**, SAL vs PSI24h $p=.6818$ |
| **5E** | | | $F(3, 15)=4.277$, **$p=.0227$** | SAL < PSI6h $p=.0525$, SAL < PSI12h **$p=.0103$**, SAL vs PSI24h $p=.3139$ |
| **5F** | | | $F(3, 15)=2.192$, $p=.1314$ | SAL vs PSI6h $p=.2117$, SAL > PSI12h $p=.0931$, SAL vs PSI24h $p=.8480$ |
| **5G** | | | $F(3, 15)=4.426$, **$p=.0203$** | SAL > PSI6h **$p=.0335$**, SAL > PSI12h **$p=.0129$**, SAL vs PSI24h $p=.4010$ |
| **5I1** | Two-way ANOVA | SAL n=3; PSI12h n=5 | Treatment $F(1, 300)=9.214$, **$p=.0026$**<br>Distance $F(49, 300)=10.34$, **$p<.0001$**<br>Interaction $F(49, 300)=0.8619$, $p=.7310$ | |
| **5I2** | Unpaired t-test | | $t(6)=2.030$, $p=.0887$ | |
| **5J1** | Two-way ANOVA; Sidak's MC | | Treatment $F(1, 300)=22.38$, **$p<.0001$**<br>Distance $F(49, 300)=3.549$, **$p<.0001$**<br>Interaction $F(49, 300)=1.459$, **$p=.0313$** | 1050 microns: SAL < PSI12h **$p=.0493$**<br>1110 microns: SAL < PSI12h **$p=.0166$** |
| **5J2** | Unpaired t-test | | $t(6)=3.102$, **$p=.0211$** | |
| **5K1** | Two-way ANOVA | | Treatment $F(1, 300)=19.16$, **$p<.0001$**<br>Distance $F(49, 300)=3.436$, **$p<.0001$**<br>Interaction $F(49, 300)=0.8322$, $p=.7794$ | |
| **5K2** | Unpaired t-test | | $t(6)=3.097$, **$p=.0212$** | |
| **5M** | Two-way ANOVA; Bonferroni's MC | Non-ABA SAL n=4; Non-ABA PSI6h n=5; ABA SAL n=5; ABA PSI n=7 | Treatment $F(1, 17)=0.5663$, $p=.4620$<br>ABA Exposure $F(1, 17)=0.4685$, $p=.5029$<br>Interaction $F(1, 17)=1.208$, $p=.2871$ | |
| **5N** | | | Treatment $F(1, 17)=15.50$, **$p=.0011$**<br>ABA Exposure $F(1, 17)=0.5340$, $p=.4749$<br>Interaction $F(1, 17)=0.02080$, $p=.8870$ | Non-ABA: SAL < PSI6h **$p=.0298$**<br>ABA: SAL < PSI **$p=.0206$** |
| **5O** | | | Treatment $F(1, 17)=9.038$, **$p=.0079$**<br>ABA Exposure $F(1, 17)=0.003860$, $p=.9512$<br>Interaction $F(1, 17)=0.5698$, $p=.4607$ | Non-ABA: SAL > PSI6h **$p=.0463$**<br>ABA: SAL vs PSI $p=.2102$ |

| | | | | |
|---|---|---|---|---|
| **5Q** | | | Treatment $F(1, 17)$=4.587, ***p*=.0470**<br>ABA Exposure $F(1, 17)$=5.098, ***p*=.0374**<br>Interaction $F(1, 17)$=15.26, ***p*=.0011** | Non-ABA: SAL vs PSI6h *p*=.5152<br>ABA: SAL > PSI ***p*=.0005** |
| **5R** | | | Treatment $F(1, 17)$=12.12, ***p*=.0029**<br>ABA Exposure $F(1, 17)$=2.048, *p*=.1705<br>Interaction $F(1, 17)$=5.606, ***p*=.0300** | Non-ABA: SAL vs PSI6h *p*=.9394<br>ABA: SAL > PSI ***p*=.0007** |

### Statistics Table - Supplementary Figure 1

| Figure | Statistical Test | Group n | Main analysis result |
|---|---|---|---|
| **S1A** | Unpaired t-test | SAL n=8; PSI n=12 | *t*(18)=2.072, *p*=.0529 |
| **S1B** | | | *t*(18)=0.5462, *p*=.5916 |
| **S1C** | | | *t*(18)=1.431, *p*=.1695 |
| **S1D** | | | *t*(18)=1.762, *p*=.0951 |
| **S1E** | | | *t*(18)=1.038, *p*=.3132 |
| **S1F** | | | *t*(18)=2.221, ***p*=.0394** |
| **S1G** | | | *t*(18)=0.2361, *p*=.8160 |
| **S1H** | | | *t*(23)=1.757, *p*=.0959 |
| **S1I** | | SAL n=15; PSI n=16 | *t*(29)=0.7545, *p*=.4566 |
| **S1J** | | | *t*(29)=0.9432, *p*=.3534 |
| **S1K** | | | *t*(29)=1.185, *p*=.2456 |
| **S1L** | | | *t*(29)=1.475, *p*=.1509 |
| **S1M** | | | *t*(29)=1.306, *p*=.2019 |
| **S1N** | | | *t*(29)=1.602, *p*=.1199 |
| **S1O** | | SAL n=14; PSI n=16 | *t*(28)=1.600, *p*=.1209 |
| **S1P** | | SAL n=15; PSI n=16 | *t*(29)=1.669, *p*=.1059 |
| **S1Q** | | | *t*(29)=1.195, *p*=.2419 |
| **S1R** | | | *t*(29)=1.120, *p*=.2721 |
| **S1S** | | | *t*(29)=1.285, *p*=.2089 |
| **S1T** | | | *t*(29)=1.141, *p*=.2631 |
| **S1U** | | SAL n=14; PSI n=16 | *t*(28)=1.341, *p*=.1907 |
| **S1V** | | SAL n=15; PSI n=16 | *t*(29)=0.4086, *p*=.6858 |
| **S1W** | | SAL n=14; PSI n=16 | *t*(28)=1.474, *p*=.1517 |

Statistics Table - Supplementary Figure 2

| Figure | Statistical Test | Group n | Main analysis result | Post-hoc multiple comparisons of interest |
|---|---|---|---|---|
| **S2C** | RM Two-way ANOVA; Bonferroni's MC | SAL n=12; PSI n=13 | Schedule $F(1, 23)=1.297$, $p=.2666$<br>Treatment $F(1, 23)=1.237$, $p=.2775$<br>Interaction $F(1, 23)=0.5375$, $p=.4709$ | |
| **S2D** | | | Schedule $F(1, 23)=79.64$, $\boldsymbol{p<.0001}$<br>Treatment $F(1, 23)=3.487$, $p=.0746$<br>Interaction $F(1, 23)=5.043$, $\boldsymbol{p=.0346}$ | PR: SAL vs PSI $p>.9999$<br>R-PR: SAL < PSI $\boldsymbol{p=.0209}$ |
| **S2E** | | | Schedule $F(1, 23)=0.03777$, $p=.8476$<br>Treatment $F(1, 23)=0.4364$, $p=.5154$<br>Interaction $F(1, 23)=0.7408$, $p=.3983$ | |
| **S2F** | | | Schedule $F(1, 23)=111.5$, $\boldsymbol{p<.0001}$<br>Treatment $F(1, 23)=1.011$, $p=.3251$<br>Interaction $F(1, 23)=0.4536$, $p=.5073$ | |
| **S2G** | | | Schedule $F(1, 23)=0.007914$, $p=.9299$<br>Treatment $F(1, 23)=9.499$, $\boldsymbol{p=.0053}$<br>Interaction $F(1, 23)=0.2956$, $p=.5919$ | PR: SAL vs PSI $p=.1324$<br>R-PR: SAL < PSI $\boldsymbol{p=.0235}$ |
| **S2H** | | | Schedule $F(1, 23)=237.0$, $\boldsymbol{p<.0001}$<br>Treatment $F(1, 23)=9.268$, $\boldsymbol{p=.0058}$<br>Interaction $F(1, 23)=5.664$, $\boldsymbol{p=.0260}$ | PR: SAL vs PSI $p>.9999$<br>R-PR: SAL < PSI $\boldsymbol{p=.0009}$ |
| **S2I** | | | Schedule $F(1, 23)=0.5233$, $p=.4767$<br>Treatment $F(1, 23)=1.091$, $p=.3072$<br>Interaction $F(1, 23)=0.2536$, $p=.6193$ | |
| **S2J** | | | Schedule $F(1, 23)=2.782$, $p=.1089$<br>Treatment $F(1, 23)=1.863$, $p=.1854$<br>Interaction $F(1, 23)=1.341$, $p=.2588$ | |
| **S2K** | | | Schedule $F(1, 23)=0.2172$, $p=.6455$<br>Treatment $F(1, 23)=1.485$, $p=.2353$<br>Interaction $F(1, 23)=0.1447$, $p=.7072$ | |
| **S2L** | | | Schedule $F(1, 23)=0.03441$, $p=.8545$<br>Treatment $F(1, 23)=0.4006$, $p=.5330$<br>Interaction $F(1, 23)=0.6415$, $p=.4314$ | |
| **S2M** | | | Schedule $F(1, 23)=1.456$, $p=.2398$ | |

| | | | |
|---|---|---|---|
| | | | Treatment *F*(1, 23)=1.892, *p*=.1822<br>Interaction *F*(1, 23)=0.3208, *p*=.5766 |

### Statistics Table - Supplementary Figure 3

| Figure | Statistical Test | Group n | Main analysis result |
|---|---|---|---|
| **S3C** | Unpaired t-test | SAL+SAL n=8; WAY+SAL n=7 | *t*(13)=3.939, ***p*=.0017** |
| **S3D** | | | *t*(13)=2.005, *p*=.0662 |
| **S3E** | | | *t*(13)=2.140, *p*=.0519 |
| **S3F** | | | *t*(13)=3.593, ***p*=.0033** |
| **S3G** | | | *t*(13)=1.471, *p*=.1650 |
| **S3H** | | | *t*(13)=0.1780, *p*=.8615 |
| **S3I** | | | *t*(13)=0.9253, *p*=.3717 |
| **S3J** | | | *t*(13)=1.314, *p*=.2114 |
| **S3K** | | | *t*(13)=0.2177, *p*=.8310 |
| **S3L** | | | *t*(13)=1.043, *p*=.3159 |
| **S3M** | | | *t*(13)=2.636, ***p*=.0206** |

### Statistics Table - Supplementary Figure 4

| Figure | Statistical Test | Group n | Main analysis result |
|---|---|---|---|
| **S4B** | Unpaired t-test | SAL+SAL n=15; KTN+SAL n=11 | *t*(24)=2.073, ***p*=.0491** |
| **S4C** | | | *t*(24)=2.465, ***p*=.0212** |
| **S4D** | | | *t*(24)=2.492, ***p*=.0200** |
| **S4E** | | | *t*(24)=3.496, ***p*=.0019** |
| **S4F** | | | *t*(24)=2.332, ***p*=.0284** |
| **S4G** | | SAL+SAL n=15; KTN+SAL n=6 | *t*(19)=2.468, ***p*=.0233** |
| **S4H** | | SAL+SAL n=14; KTN+SAL n=2 | *t*(14)=-.1394, *p*=.8912 |
| **S4I** | | SAL+SAL n=15; KTN+SAL n=11 | *t*(24)=3.509, ***p*=.0018** |
| **S4K** | | SAL+PSI n=16; KTN+PSI n=12 | *t*(26)=1.878, *p*=.0716 |
| **S4L** | | | *t*(26)=2.165, ***p*=.0397** |
| **S4M** | | | *t*(26)=2.287, ***p*=.0306** |
| **S4N** | | | *t*(26)=3.085, ***p*=.0048** |
| **S4O** | | | *t*(26)=2.832, ***p*=.0088** |
| **S4P** | | SAL+PSI n=16; KTN+PSI n=11 | *t*(25)=3.158, ***p*=.0041** |
| **S4Q** | | SAL+PSI n=16; KTN+PSI n=8 | *t*(22)=1.652, *p*=.1128 |
| **S4R** | | SAL+PSI n=16; KTN+PSI n=12 | *t*(26)=1.449, *p*=.1592 |

### Statistics Table - Supplementary Figure 5

| Figure | Statistical Test | Group n | Main analysis result | Post-hoc multiple comparisons of interest |
|---|---|---|---|---|
| **S5A** | One-way ANOVA; Dunnett's MC | SAL n=4; PSI6h n=5; PSI12h n=5; PSI24h n=5 | *F*(3, 15)=0.5145, *p*=.6785 | |
| **S5B** | | | *F*(3, 15)=3.870, ***p*=.0312** | SAL < PSI6h *p*=.0867, SAL < PSI12h ***p*=.0137**; SAL vs PSI24h *p*=.4043 |
| **S5C** | | | *F*(3, 15)=1.825, *p*=.1859 | |
| **S5D** | | | *F*(3, 15)=2.286, *p*=.1204 | |
| **S5E** | | | *F*(3, 15)=3.429, ***p*=.0445** | SAL > PSI6h *p*=.0760, SAL > PSI12h ***p*=.0295**; SAL vs PSI24h *p*=.6046 |
| **S5F** | | | *F*(3, 15)=0.9926, *p*=.4230 | |
| **S5G** | | | *F*(3, 15)=0.7283, *p*=.5509 | |
| **S5H** | Two-way ANOVA; Bonferroni's MC | Non-ABA SAL n=4; Non-ABA PSI n=5; ABA SAL n=5; ABA PSI n=7 | Treatment *F*(1, 17)=0.5179, *p*=.4815<br>ABA Exposure *F*(1, 17)=0.6851, *p*=.4193<br>Interaction *F*(1, 17)=0.3453, *p*=.5645 | |
| **S5I** | | | Treatment *F*(1, 17)=16.33, ***p*=.0008**<br>ABA Exposure *F*(1, 17)=0.1060, *p*=.7487<br>Interaction *F*(1, 17)=1.165, *p*=.2954 | Non-ABA: SAL < PSI ***p*=.0068**<br>ABA: SAL < PSI *p*=.0756 |
| **S5J** | | | Treatment *F*(1, 17)=2.677, *p*=.1202<br>ABA Exposure *F*(1, 17)=2.966, *p*=.2032<br>Interaction *F*(1, 17)=5.595, ***p*=.0302** | Non-ABA: SAL < PSI ***p*=.0333**<br>ABA: SAL vs PSI *p*>.9999 |
| **S5K** | | | Treatment *F*(1, 17)=4.316, *p*=.0532<br>ABA Exposure *F*(1, 17)=1.117, *p*=.3954<br>Interaction *F*(1, 17)=2925, *p*=.1054 | Non-ABA: SAL < PSI ***p*=.0446**<br>ABA: SAL vs PSI *p*>.9999 |
| **S5L** | | | Treatment *F*(1, 17)=10.58, ***p*=.0047**<br>ABA Exposure *F*(1, 17)=0.09476, *p*=.7620<br>Interaction *F*(1, 17)=1.262, *p*=.2769 | Non-ABA: SAL > PSI ***p*=.0197**<br>ABA: SAL vs PSI *p*=.2476 |
| **S5M** | | | Treatment *F*(1, 17)=4.177, *p*=.0568<br>ABA Exposure *F*(1, 17)=4.645, ***p*=.0458**<br>Interaction *F*(1, 17)=10.37, ***p*=.0050** | Non-ABA: SAL vs PSI *p*=.8914<br>ABA: SAL > PSI ***p*=.0018** |
| **S5N** | | | Treatment *F*(1, 17)=10.91, ***p*=.0042**<br>ABA Exposure *F*(1, 17)=0.7079, *p*=.4118<br>Interaction *F*(1, 17)=2.092, *p*=.1663 | Non-ABA: SAL vs PSI *p*=.4691<br>ABA: SAL > PSI ***p*=.0043** |
| **S5O** | | | Treatment *F*(1, 17)=0.4758, *p*=.4996 | |

| | | | | |
|---|---|---|---|---|
| | | | ABA Exposure *F*(1, 17)=0.2064, *p*=.6553<br>Interaction *F*(1, 17)=2.163, *p*=.1597 | |
| **S5P** | | | Treatment *F*(1, 17)=12.12, ***p*=.0029**<br>ABA Exposure *F*(1, 17)=1.177, *p*=.2930<br>Interaction *F*(1, 17)=0.6974, *p*=.4153 | Non-ABA: SAL vs PSI *p*=.1941<br>ABA: SAL < PSI ***p*=.0088** |
| **S5Q** | | | Treatment *F*(1, 17)=1.148, *p*=.2898<br>ABA Exposure *F*(1, 17)=1.530, *p*=.2329<br>Interaction *F*(1, 17)=5.336, ***p*=.0337** | Non-ABA: SAL < PSI *p*=.0768<br>ABA: SAL vs PSI *p*=.7193 |
| **S5R** | | | Treatment *F*(1, 17)=2.832, *p*=.1107<br>ABA Exposure *F*(1, 17)=0.4715, *p*=.5016<br>Interaction *F*(1, 17)=2.566, *p*=.1276 | Non-ABA: SAL < PSI *p*=.0872<br>ABA: SAL vs PSI *p*>.9999 |
| **S5S** | | | Treatment *F*(1, 17)=6.732, ***p*=.0189**<br>ABA Exposure *F*(1, 17)=0.1640, *p*=.6905<br>Interaction *F*(1, 17)=0.1327, *p*=.7201 | Non-ABA: SAL vs PSI *p*=.1322<br>ABA: SAL vs PSI *p*=.2163 |
| **S5T** | | | Treatment *F*(1, 17)=3.792, *p*=.0682<br>ABA Exposure *F*(1, 17)=4.213, *p*=.0558<br>Interaction *F*(1, 17)=15.16, ***p*=.0012** | Non-ABA: SAL vs PSI *p*=.4277<br>ABA: SAL > PSI ***p*=.0007** |
| **S5U** | | | Treatment *F*(1, 17)=10.01, ***p*=.0057**<br>ABA Exposure *F*(1, 17)=2.859, *p*=.1091<br>Interaction *F*(1, 17)=7.599, ***p*=.0135** | Non-ABA: SAL vs PSI *p*>.9999<br>ABA: SAL > PSI ***p*=.0006** |

### Statistics Table - Supplementary Figure 6

| Figure | Statistical Test | Group n | Main analysis result | Post-hoc multiple comparisons of interest |
|---|---|---|---|---|
| **S6A** | RM Two-way ANOVA; Mixed-effects analysis; Bonferroni's MC | PSI-S n=11; PSI-R n=8 | Day *F*(1, 17)=14.78, ***p*<.0001**<br>ABA Outcome *F*(1, 17)=6.341, ***p*=.0221**<br>Interaction *F*(1, 17)=3.196, ***p*=.0071** | PSI-S > PSI-R Day 7 *p*=.0697 |
| **S6B** | | | Day *F*(1, 17)=27.22, ***p*<.0001**<br>ABA Outcome *F*1, 17)=13.98, ***p*=.0016**<br>Interaction *F*(1, 17)=7.957, ***p*<.0001** | PSI-S > PSI-R Day 1 ***p*=.0354;** PSI-S > PSI-R Day 3 ***p*=.0173**; PSI-S > PSI-R Day 4 ***p*=.0029;** PSI-S > PSI-R Day 5 ***p*=.0183** |

### Statistics Table - Supplementary Figure 7

| Figure | Statistical Test | Group n | Main analysis result |
|---|---|---|---|
| **S7E** | Unpaired t-test | SAL (n=11) vs PSI (n=12) | *t*(21)=0.4179, *p*=.6802 |
| **S7F** | | | *t*(21)=0.9673, *p*=.3444 |
| **S7G** | | | Treatment *F*(1, 21)=0.4156, *p*=.5561<br>Phase *F*(1, 21)=169.4, ***p*<.0001**<br>Interaction *F*(1, 21)=0.3644, *p*=.5526 |
